# Genetic Architecture of Maize Rind Strength Revealed by the Analysis of Divergently Selected Populations

**DOI:** 10.1101/2020.04.14.041517

**Authors:** Rohit Kumar, Abiskar Gyawali, Ginnie D. Morrison, Christopher A. Saski, Daniel J. Robertson, Nishanth Tharayil, Robert J. Schaefer, Timothy M. Beissinger, Rajandeep S. Sekhon

## Abstract

Stalk lodging, breakage of the stalk at or below the ear, causes substantial yield losses in maize. The strength of the stalk rind, commonly measured as rind penetrometer resistance (RPR), is an important contributor to stalk lodging resistance. To enhance RPR genetic architecture, we conducted selection mapping on populations developed by 15 cycles of divergent selection for high (C15-H) and low (C15-L) RPR. We also performed time-course transcriptome and metabolic analyses on developing stalks of high (Hrpr1) and low (Lrpr1) RPR inbred lines derived from the C15-H and C15-L populations, respectively. Divergent selection significantly altered allele frequencies at 3,656 and 3,412 single nucleotide polymorphisms (SNP) in the C15-H and C15-L populations, respectively. While the majority of the SNPs under selection were unique, 110 SNPs were common in both populations indicating the fixation of alleles with alternative effects. Remarkably, preferential selection on the genomic regions associated with lignin and polysaccharide biosynthesis genes was observed in C15-H and C15-L populations, respectively. This observation was supported by higher lignification and lower extractability of cell wall-bound sugars in Hrpr1 compared to Lrpr1. Tricin, a monolignol important for incorporation of lignin in grass cell walls, emerged as a key determinant of the different cell wall properties of Hrpr1 and Lrpr1. Integration of selection mapping with transcriptomics and previous genetic studies on RPR identified 40 novel candidate genes including Zm*MYB31, ZmNAC25, ZmMADS1*, two *PAL* paralogues, two lichenases, *ZmEXPA2, ZmIAA41*, and *Caleosin*. Enhanced mechanistic and genetic understanding of RPR provides a foundation for improved stalk lodging resistance.

## Introduction

Maize (*Zea mays* L.) is one of the most important cereal crops in the world. Stalk lodging, which refers to the breakage of the stalk at or below the ear, is estimated to cause 5% - 25% annual yield losses in maize (Zuber and Kang, 1978; Flint-Garcia et al., 2003). Furthermore, stalk lodging often results in decreased grain quality and an increased presence of pests and disease due to the decay of fallen (i.e., lodged) grains. Increased use of nitrogen and higher planting density associated with the cultivation of higher-yielding maize hybrids are poised to further increase stalk lodging incidence (Yang et al., 2019). Improving the extractability of sugars for forage or biofuels also accompanies decreased stalk strength and increased lodging (Pedersen et al., 2005; Feltus and Vandenbrink, 2012). Finally, ever-worsening climate and associated extreme weather events are expected to enhance yield losses associated with lodging. A comprehensive understanding of the genetic architecture of stalk strength is key to the successful breeding and biotechnological intervention targeted to improve grain yield and biomass quality.

Phenotypic assessment of stalk lodging resistance has been challenging primarily because of difficulties in determining the aspects of stalk strength that translate into higher lodging resistance in field conditions. Indirect methods for prediction of stalk lodging resistance include counting lodged plants in field evaluations (Robertson et al., 2016), analyzing the chemical composition of stalks (Davidson and Phillips, 1930; Appenzeller et al., 2004; Ching et al., 2006), measuring stalk bending strength (Robertson et al., 2014; Robertson et al., 2016), stalk crushing strength (Zuber and Grogan, 1961; Undersander et al., 1977; Zuber et al., 1980), and rind penetrometer resistance (Zuber et al., 1980; Sibale et al., 1992). Rind strength is proposed to contribute to 50–80% of stalk strength (Zuber et al., 1980). Rind strength can be assessed by forcing a small probe through a plant stalk and measuring the maximum strength required to puncture the rind. This method, which measures rind penetrometer resistance (RPR), also known as rind puncture resistance, has been used throughout most of the 20th century to investigate stalk strength (Khanna, 1935). RPR has been shown to be significantly and negatively correlated with naturally occurring stalk lodging in the field (Dudley, 1994; Kang et al., 1999; Jampatong et al., 2000; Hu et al., 2012; Sekhon et al., 2019). In a recent and comprehensive study, RPR data collected on 47 maize hybrids in three environments was shown to be a reliable predictor of stalk lodging incidence of these hybrids observed in 98 temporally and spatially distinct environments (Sekhon et al., 2019). To summarize, RPR offers a high-throughput phenotyping method for artificial selection in breeding programs and for genetic studies aimed to understand the genetic architecture of lodging resistance.

Several genetics studies have reported RPR as a quantitative trait and identified quantitative trait loci (QTL) or single nucleotide polymorphisms (SNP) putatively associated with this trait (Heredia et al., 1996; Flint-Garcia et al., 2003; Hu et al., 2012; Peiffer et al., 2013). To exploit RPR for improvement of stalk lodging resistance, a divergent (i.e., bidirectional) selection experiment in Missouri Second Cycle Stiff Stalk Synthetic (MoSCSSS), a yellow dent synthetic population formed from intermating of 14 inbred lines, was initiated using recurrent S_0_ selection. While the original MoSCSSS population, designated as cycle 0 (C0), had RPR value of 3.55 kgf, twelve cycles of selection for high and low RPR from this population resulted in the development of high and low RPR populations with RPR value of 8.52 kgf and 2.0 kgf, respectively (Martin et al., 2004). Multiple rounds of such divergent selection have the potential to fix alternative alleles with contrasting effects on RPR or to substantially alter the allele frequencies of alternate alleles in these populations. Therefore, the populations developed by divergent selection were for the discovery of QTL underlying RPR (Flint-Garcia et al., 2003). To this end, F_2:3_ families obtained from the high RPR population derived from 10 cycles of selection were crossed with: 1) F_2:3_ families obtained from the low RPR population derived from 11 cycles of selection, 2) an S_1_ plant with low RPR selected from an unrelated population (MoSQB-Low), and 3) with an inbred line Mo47. The resulting mapping populations were used to identify 26 QTL for RPR (Flint-Garcia et al., 2003). Evaluation of recombinant inbred lines (RILs) from a maize nested association mapping population and a large collection of diverse inbred lines identified several significant single nucleotide polymorphisms (SNPs) and QTL associated with RPR (Peiffer et al., 2013). Evaluation of RILs derived from B73 and a high kernel oil inbred resulted in the identification of nine QTL for RPR (Hu et al., 2012), and evaluation of two sets of RILs reported seven QTL (Li et al., 2014). These studies provide rich information about genomic regions that control RPR, and identification of the underlying genes will help enhance stalk strength. Recently, cloning of a major QTL underlying RPR resulted in the identification of a novel gene *stiff1,* and knockdown of *stiff1*, either naturally through a transposon insertion in the gene promoter or experimentally through gene editing, enhanced stalk strength (Zhang et al., 2020). However, progress in cloning and characterizing the underlying genes has been slow and requires novel genomic approaches to improve the resolution of existing genetic information on the genomic regions governing RPR.

Divergent selection experiments enable a direct assessment of the response of a genome to selection for a trait. This in turn leads to better understanding and a greater ability to characterize the genetic architecture of the given trait. Populations derived from divergent selection are an example of experimental evolution (Turner et al., 2011), and hence exhibit differences in allele frequency between pre- and post-selection populations. Modern genotyping approaches allow the assessment of allele frequency changes in experimentally-evolved populations to map putatively causal loci (Hirsch et al., 2014; Lorenz et al., 2015; Kelly and Hughes, 2019). The field of gene-mapping has been dominated by phenotype-to-genotype correlation analyses including genome-wide association (GWA) and QTL mapping for several decades (Pascual et al., 2016; Tam et al., 2019). Scanning for selective sweeps is an additional tactic that can be effectively utilized to identify trait-relevant genomic regions and genes in selected populations (Nielsen et al., 2005; Kimura et al., 2007; Qanbari et al., 2012; Beissinger et al., 2014; Chen et al., 2018; Grainger et al., 2018; You et al., 2018). For agricultural species, previous breeding and long-term selection studies have already generated many such populations (Olsen and Wendel, 2013). In contrast to GWA and QTL mapping studies, selection scans do not depend on phenotypes from genotyped individuals. Instead, with the given knowledge that selection for a specific trait or traits has occurred, a scan for selected loci may proceed without any additional phenotyping (Gerke et al., 2015). In addition to eliminating time-intensive phenotyping, selection mapping can serve as an independent mapping technique to corroborate region-trait associations identified via other methods.

The current study presents phenotypic, genomic, and transcriptomic analyses of divergently selected maize populations to gain mechanistic insights into RPR in maize. Specifically, we 1) examined the phenotypic changes associated with divergent selection in the population obtained from additional cycles of selections than those reported earlier, and inbred lines developed from these populations, 2) examined the selection signatures to identify the divergently selected genomic segments in the populations, 3) performed transcriptomic and metabolic analyses on the inbred lines with high and low RPR derived from the aforementioned divergently selected populations, and 4) combined the data from the three experiments and previously published studies to begin constructing a comprehensive picture of the genetic architecture of RPR in maize. Results from this study will boost efforts towards the identification of genes and genetic elements underlying RPR and alleviation of lodging-related losses in cereals.

## Results

### Divergent selection resulted in significant changes in rind penetrometer resistance

Phenotypic changes after twelve cycles of divergent selection for RPR on MoSCSSS (C0) population have already been reported (Martin et al., 2004). To test the effect of three additional cycles of selection in each direction, we recorded RPR in C15-H and C15-L populations and found significant differences among the three populations (Figure 1). Mean RPR of C0, C15-L, and C15-H was 4.55 Kgf, 2.1 Kgf, and 11.9 kgf, respectively.

**Figure 1.**
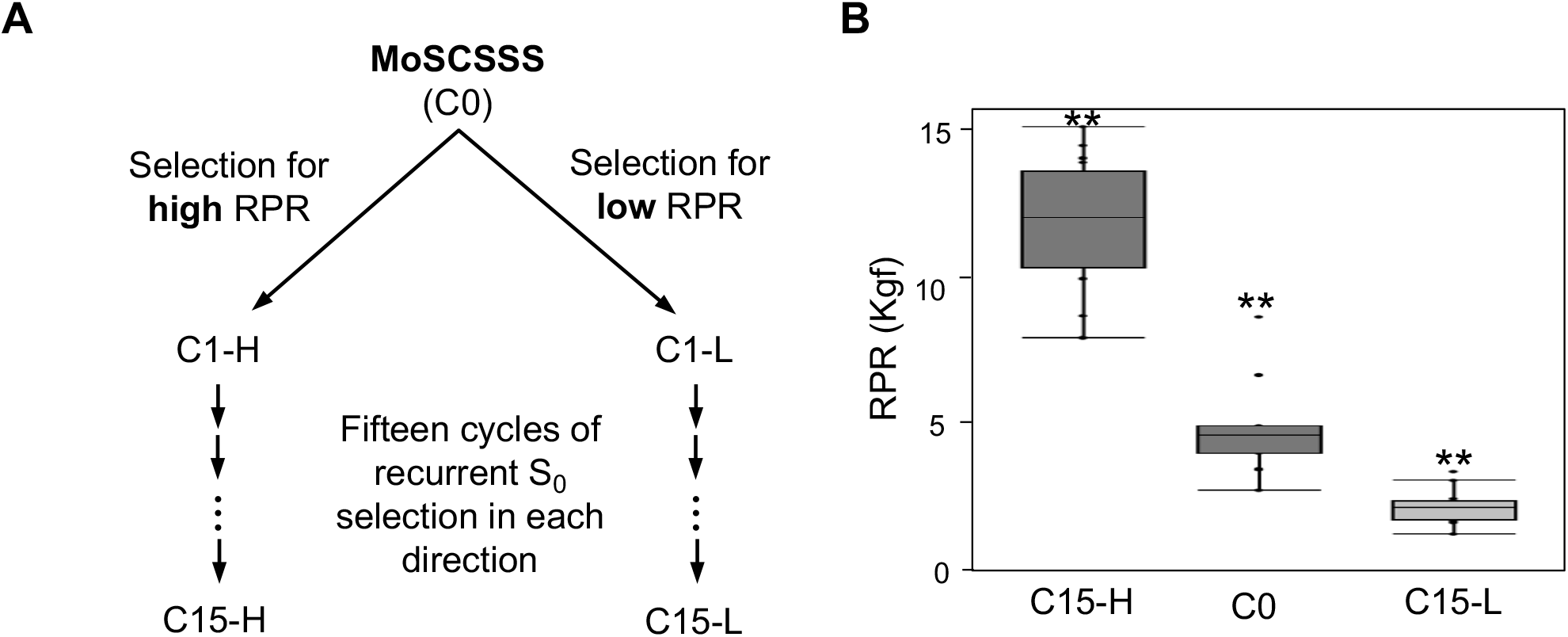
Divergent selection for rind penetrometer resistance (RPR) and the associated phenotypic changes. A. Schema for 15 cycles of divergent selection performed by Dr. Larry Darrah and colleagues. B. Box plots showing phenotypic divergence of the two populations from the C0 population. Populations are shown on the *x*-axis and RPR in kilograms of force (Kgf) is shown on the *y*-axis. Two asterisks indicate significance at *p* < 0.01.

### Divergent selection significantly modified the frequency of several SNPs genome-wide

We computed *F_ST_* and changes in allele frequency between C0 and C15-L, and between C0 and C15-H, as well as *F_ST_* values for these comparisons (Figure 2). Highly significant correlations between changes in allele frequency and *F_ST_* were observed both for high RPR selection (*r* = 0.9644; *p* < 1e-324) and low RPR selection (*r* = 0.9641; *p* < 1e-324). Using allele frequency change from C0 to C15-H to establish significance thresholds, we identified 3,656 significant SNPs that were putatively selected for high RPR (Figure 2, Table S1). Likewise, based on allele frequency change from C0 to C15-L, we identified 3,412 SNPs that showed significant evidence of selection for low RPR (Figure 2, Table S1). In both cases, observed allele frequency changes cannot be explained by drift alone, although we cannot exclude the possibility that other nonneutral force(s) drove changes in allele frequency. Those forces could include unintentional selection on additional traits besides high RPR and low RPR, such as general fitness. Of these significant SNPs, 110 SNPs were selected in both directions (Figure 2, Table S1) suggesting fixation of alternative alleles of certain genes. Bootstrap resampling indicated that the extent of such overlap is significant (*p* ≤ 1e-4). For selection in both directions, there were clusters of many SNPs showing significant selection signals. Such clusters included the distal-arm of chromosome 2 for C15-H and the distal-arm of chromosome 7 for C15-L, respectively (Figure 2). Regions represented by these clusters likely harbor large-effect genes that experienced strong selection during early generations of the experiment.

**Figure 2.**
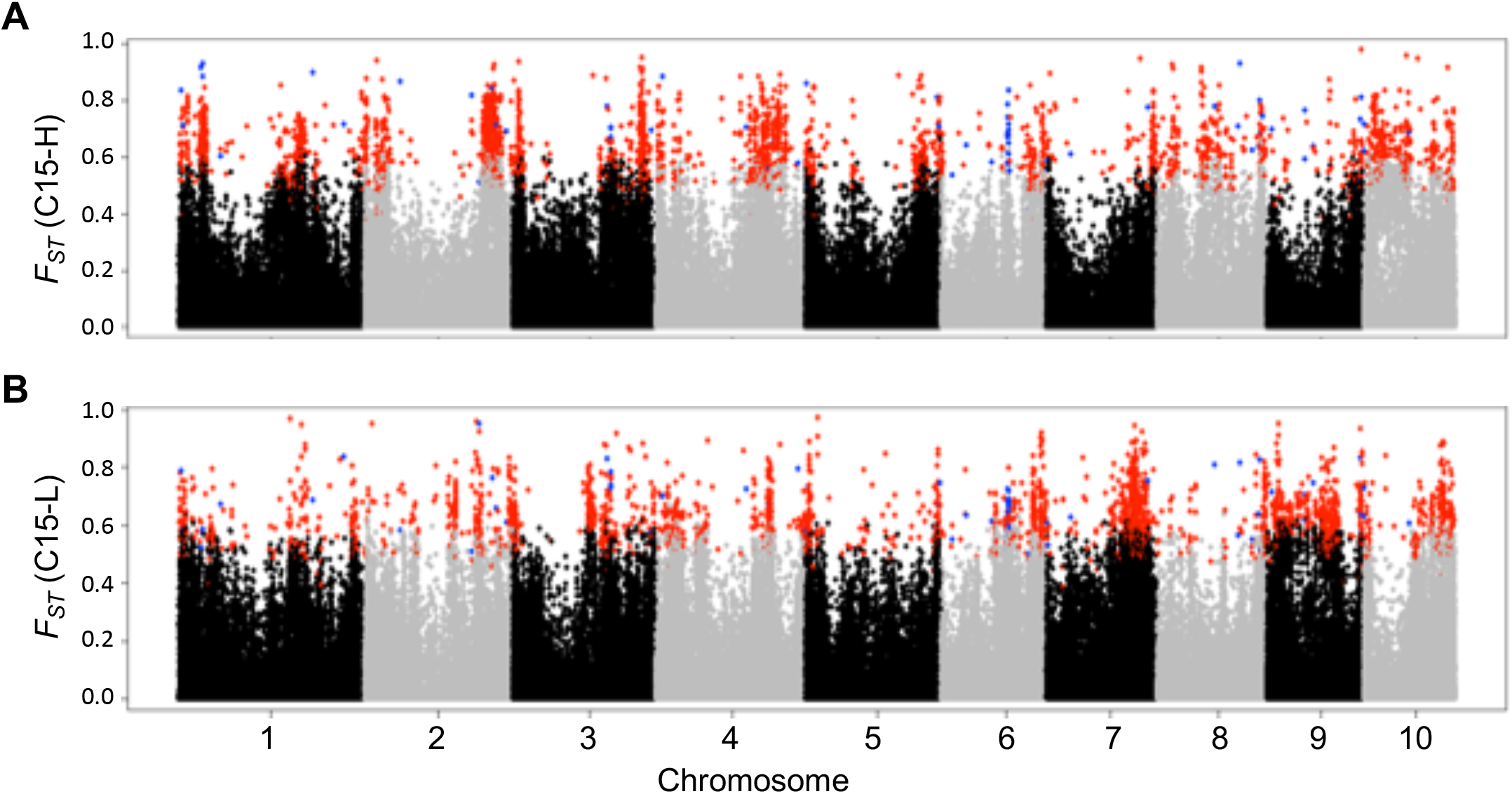
Identification of SNPs associated with divergent selection. Shown here are plots of significantly selected SNPs obtained from comparison between C15-H and C0 (A), and between C15-L and C0 (B). Each dot represents a SNP, vertical position of the dot represents *F_ST_* value shown on *y*-axis, and the horizontal location of the dot represents physical location of the SNP on a chromosome shown on *x*-axis. Grey and black color differentiates the chromosomes. Red and blue colored dots represent SNPs that were significant according to our simulation strategy. Blue colored dots represent significant SNPs that are common in both high (C15-H) and low (C15-L) populations.

### Divergence in RPR phenotype accompanied with distinct chemical composition of internodes

C15-H and C15-L populations are maintained in a heterozygous state (Flint-Garcia et al., 2003; Martin et al., 2004). To fix and study a subset of alleles that underlie the diverged phenotype and likely regulate RPR, we developed Hrpr1 and Lrpr1 inbred lines from C15-H and C15-L populations, respectively. The RPR phenotype and transcriptome associated with RPR was measured on the 12^th^ internode which, on average, represented the internode below the ear bearing node in the inbred lines. The internodes were immature and completely lacked RPR at 0 DAS stage as indicated by a lack of detected RPR value by the rind penetrometer (Figure 3A). The two inbred lines started to significantly diverge at 6 DAS, and 9 DAS appeared to be a distinctive stage with a sharp increase in RPR of Hrpr1 compared to Lrpr1. While RPR continued to increase in both inbreds after 12 DAS, albeit at a lower rate, Hrpr1 maintained significantly higher RPR compared to Lrpr1.

**Figure 3.**
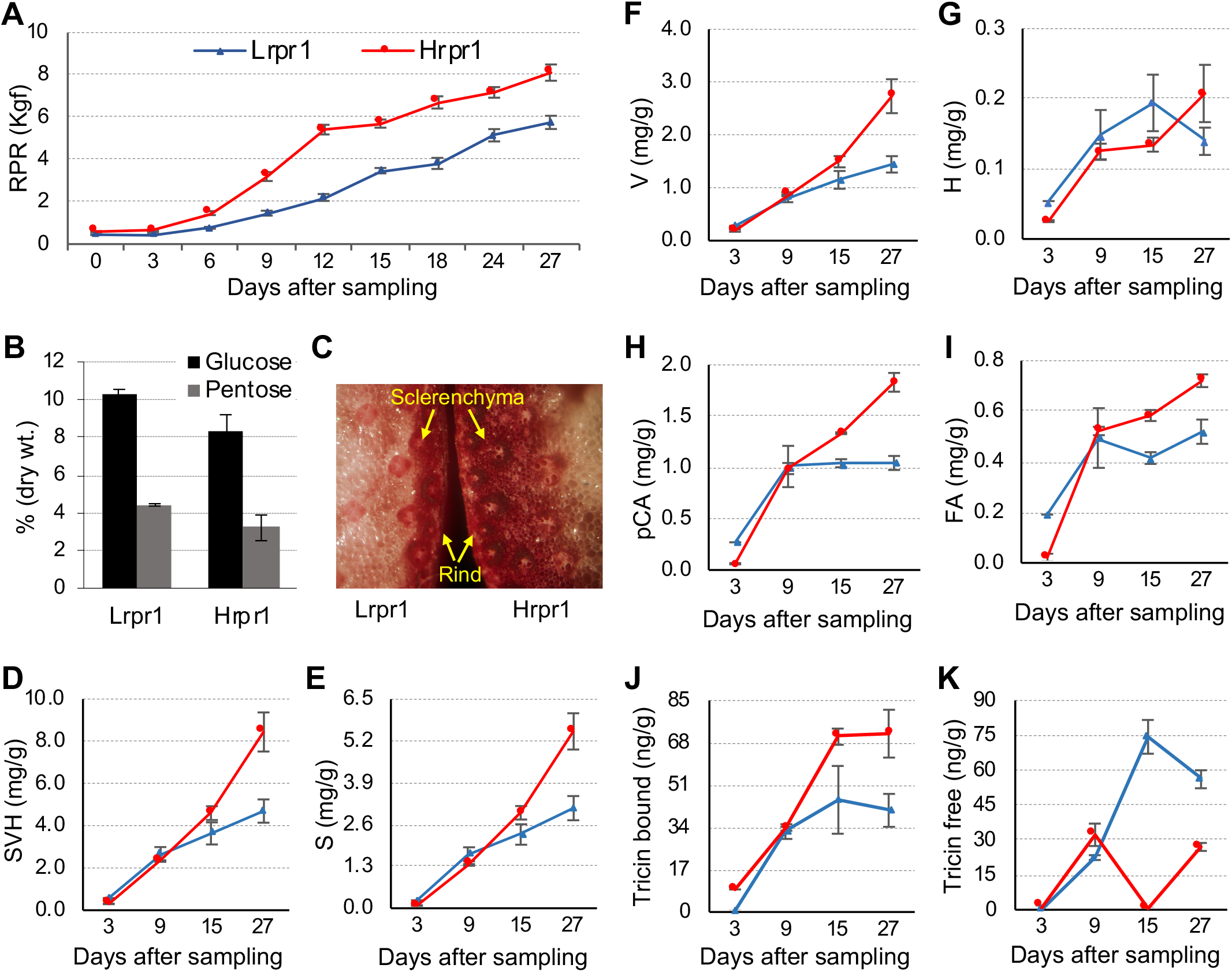
Changes in phenotype and chemical composition of internodes associated with divergent selection for rind penetrometer resistance. (A) Time course analysis showing divergence of RPR in the Hrpr1 and Lrpr1 lines. The *x*-axis shows days after sampling and the *y*-axis denotes RPR in kilograms of force (Kgf) measured by the rind penetrometer. (B) Glucose and pentose accumulation in internode of Lrpr1 and Hrpr1 inbreds. (C) Histochemical staining with Phloroglucinol-HCl of stem section of Lrpr1 (left) and Hrpr1 (right). (D-G) Accumulation of lignin monomers syringic acid (S), vanillic acid (V), and p-hydroxy benzoate (H) in internodes of Lrpr1 and Hrpr1 inbreds. Shown here is time course accumulation of SVH (D), S (E), V (F) and, H (G). (H-K) Accumulation of phenolics involved in polymerization of lignin monomers in the cell wall. Shown here is time course accumulation of p-Coumaric acid (H) and ferulic acid (I), lignin bound tricin (J), and unbound tricin (K) in the internodes of Lrpr1 and Hrpr1 inbreds.

The constituents of cell wall, including polysaccharides (cellulose and hemicellulose) and secondary metabolites (primarily lignin), are proposed to be important determinants of stalk strength (Bosch et al., 2011; Peiffer et al., 2013). To assess their role in the observed differences in RPR, we examined the accumulation of these metabolites in the internodes of Hrpr1 and Lrpr1. Remarkably, RPR levels were negatively correlated with the amount of these monosaccharides released from the internode cell wall, and significantly higher glucose and pentose was obtained from Lrpr1 compared to Hrpr1 (Figure 3B). Histological staining of the internode sections revealed higher lignification of sclerenchyma and rind-region parenchyma in Hrpr1 compared to Lrpr1 (Figure 3C). The differences in histological staining, and the release of sugars from the lignocellulosic biomass (i.e., digestibility), in grasses are attributed to increased lignification (Akin, 1989). Lignin is primarily composed of syringyl, guaiacyl, and, to a much lesser extent, p-hydroxyphenyl monomers (Halpin, 2019; Ralph et al., 2019). Lignin extraction in our study, performed in oxidizing conditions, measured syringic acid (S), vanillic acid (V), and p-hydroxybenzoate (H) (Wang et al., 2015). The total amount of these three monolignols was comparable up to 9 DAS stage in both inbred lines, but the Hrpr1 internodes accumulated higher amounts at the later stages (Figure 3D). Accumulation of S and V lignin followed the same trend (Figure 3E-F, Figure S1A-F). Accumulation of H lignin was quite low in both inbreds albeit Lrpr1 internode accumulated slightly but significantly higher amounts of this monomer compared to Hrpr1 at early stages, but this trend was reversed later in development (Figure 3G, Figure S1G-I).

Incorporation of lignin monomers in the cell wall is of paramount importance for imparting mechanical strength (Tobimatsu and Schuetz, 2019). Polymerization of lignin monomers in grass cell walls is achieved by the activity of ferulic acid (FA) and *p*-coumaric acid (pCA) (Hatfield et al., 2017). Interestingly, both pCA and FA were slightly higher in Lrpr1 at 3 DAS but eventually increased to significantly higher amounts in Hrpr1 (Figure 3H-I). Tricin is an important monolignol involved in lignification by acting as a nucleation site for polymerization of lignin monomers in grass cell walls (Lan et al., 2015). The cell wall-bound fraction of tricin, which signifies the tricin incorporated in the lignin polymer, was slightly higher in Hrpr1 at 3 DAS and, while the differences diminished at 9 DAS, Hrpr1 internodes had higher amounts in the later stages of development (Figure 3J). Conversely, the unbound fraction of tricin showed a reverse trend with a higher accumulation of Lrpr1 (Figure 3K). In summary, while the differences in lignification were minimal at the early stages of development, higher lignification of internode cell walls of Hrpr1 as compared to Lrpr1 was evident at the later stages of development.

### Divergent selection for high and low RPR resulted in enrichment of distinct pathways of cell wall synthesis

Since lignin and polysaccharides emerged as two key classes of metabolites associated with distinct RPR phenotypes in the inbred lines, we examined the effect of divergent selection on genes related to the synthesis of these metabolites. We first identified the genes and gene families associated with the synthesis of cellulose, hemicellulose, and lignin in the maize genome. This analysis identified 504 genes associated with cellulose and hemicellulose synthesis, and 341 genes associated with lignin synthesis (Table 1, Table S2). We then assessed if the significant SNPs associated with these genes were enriched in SNPs identified through selection mapping (Table S1), during divergent selection for high and low RPR. In the genome space covered by the lignin pathway genes, 45 SNPs showed a significant signal of selection in C15-H population compared to 26 SNPs the C15-L population. Bootstrap analysis further showed that C15-H population was significantly enriched for SNPs linked to the lignin pathway genes compared to whole-genome background levels of selection (*p* ≤ 0.004) while C15-L population did not show enrichment (*p* ≤ 0.491) (Figure 4). In contrast, the SNPs in the genome space covering the polysaccharide synthesis genes showed an opposite pattern. Eighty SNPs showed a significant signal of selection in the C15-L population compared to 60 enriched SNPs in the C15-H population. Bootstrap analysis indicated that the C15-L population was enriched for significant SNPs in polysaccharide genome space (*p* ≤ 0.001) while the C15-H population was not enriched (*p* ≤ 0.34) (Figure 4).

**Figure 4.**
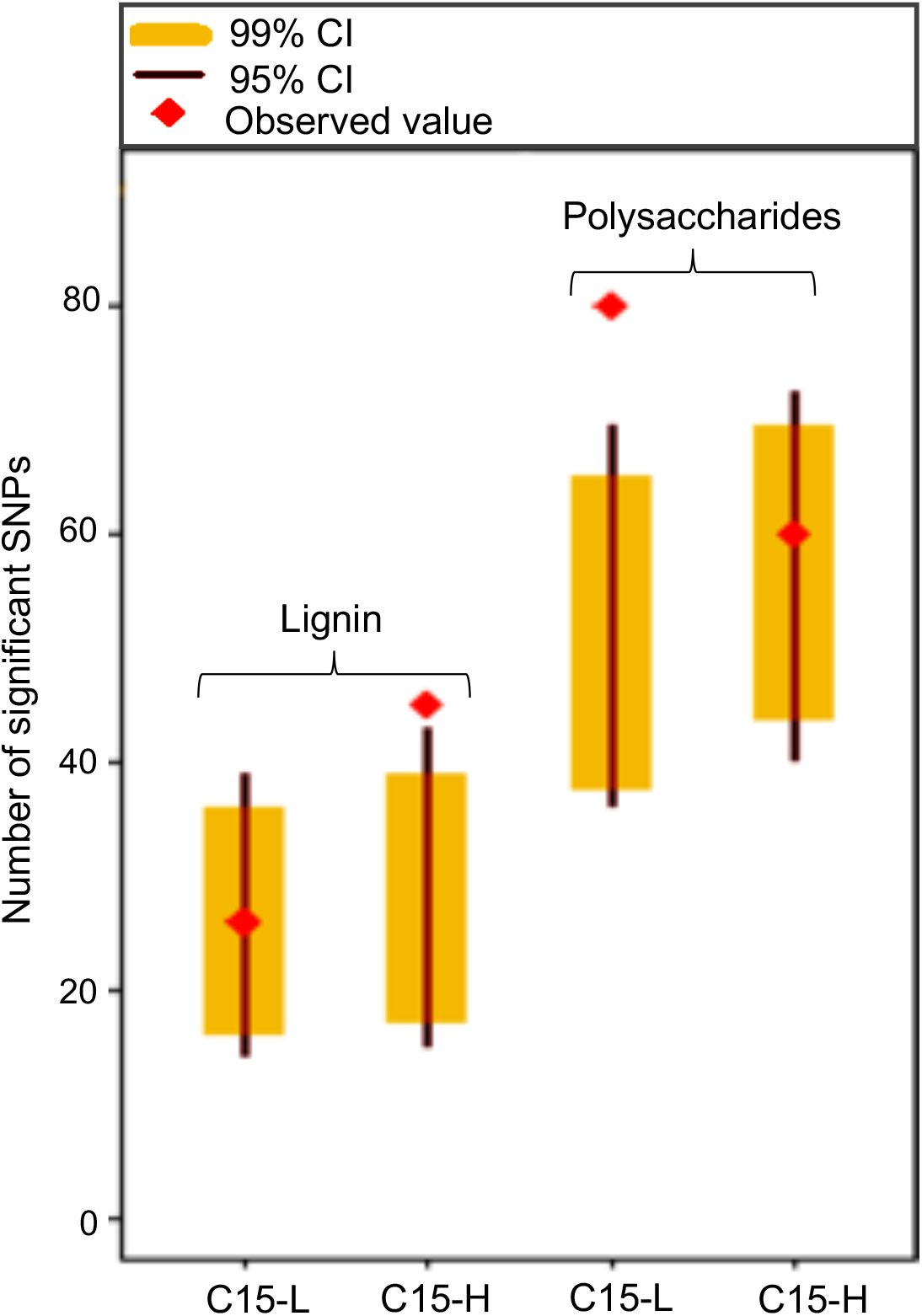
Selection of cell wall associated SNPs in low RPR (C15-L) and high RPR (C15-H) populations. Red diamonds indicate the observed value of enriched SNPs for lignin (left) and polysaccharide (right) associated genes.

**Table 1.** Identification of maize cell wall synthesis gene families.

| Cell wall component/process | Gene family | Number of genes |
| --- | --- | --- |
| Polysaccharides biosynthesis (cellulose and hemicellulose) | *Cellulose synthase (CesA/Csl)* superfamily | 83 |
|  | *GT8* superfamily (galacturonosyl transferases) | 57 |
|  | *GT47* superfamily (galactosyl, arabinosyl, xylosyl, and glucuronosyl transferases) | 54 |
|  | *GT31* superfamily (galactosyl transferases) | 32 |
| Cell wall assembly and rearrangement | *Expansin* gene family | 95 |
|  | Xyloglucan endotransglucosylase/hydrolase (*XTH*) superfamily | 35 |
| Cell wall modification and disassembly | *Polygalacturonase/ Pectin lyase (PGases)* gene family | 53 |
|  | *Pectinesterase (PME)* gene family | 48 |
| Nucleotide-sugar synthesis and interconversion | *UDP-Glc dehydrogenase (UGD)*, *UDP-GlcA decarboxylase*, *UDP-Glc 4-epimerases (UGE)*, *UDP-Xyl 4-epimerases (UXE)*, *4,6-dehydratase (GMD)* | 47 |
| Lignin biosynthesis | *Phenyl ammonia lyase (PAL)* gene family | 18 |
|  | *Caffeic acid o-methyltransferase (COMT)* gene family | 32 |
|  | *Caffeoyl-CoA O-methyltransferase (CCoAOMT)* gene family | 5 |
|  | *4-Coumaric acid CoA-ligase (4CL)* gene family | 17 |
|  | *Cinnamoyl-CoA reductases (CCR)* gene family | 36 |
|  | *p-Coumaroyl-shikimate/quinic 3'-hydroxylase (C3'H)*, *Cinnamate 4-hydroxylase (C4H)*, *Ferulate 5-hydroxylase (F5H)* gene family | 47 |
|  | *Cinnamyl alcohol dehydrogenase (CAD)* gene family | 8 |
|  | *Hydroxycinnamoyl-CoA transferase (HCT)* gene family | 42 |
|  | *Peroxidase* gene family | 114 |
|  | *Laccase* gene family | 22 |
| Total genes |  | 845 |

### Characterization of transcriptome underlying RPR

To understand the transcriptome dynamics underlying the divergently selected phenotypes for low and high RPR, we performed RNA-seq on the developing internodes of Lrpr1 and Hrpr1 at key developmental stages (Figure 5). A snapshot of transcriptome dynamics obtained by principal component analysis (Figure 5A) was consistent with developmental divergence in the RPR phenotype in Lrpr1 and Hrpr1 (Figure 3A). Both inbred lines showed close clustering at 0 and 3 DAS but dispersed considerably in the later stages indicating distinct transcriptomes of the internodes.

**Figure 5.**
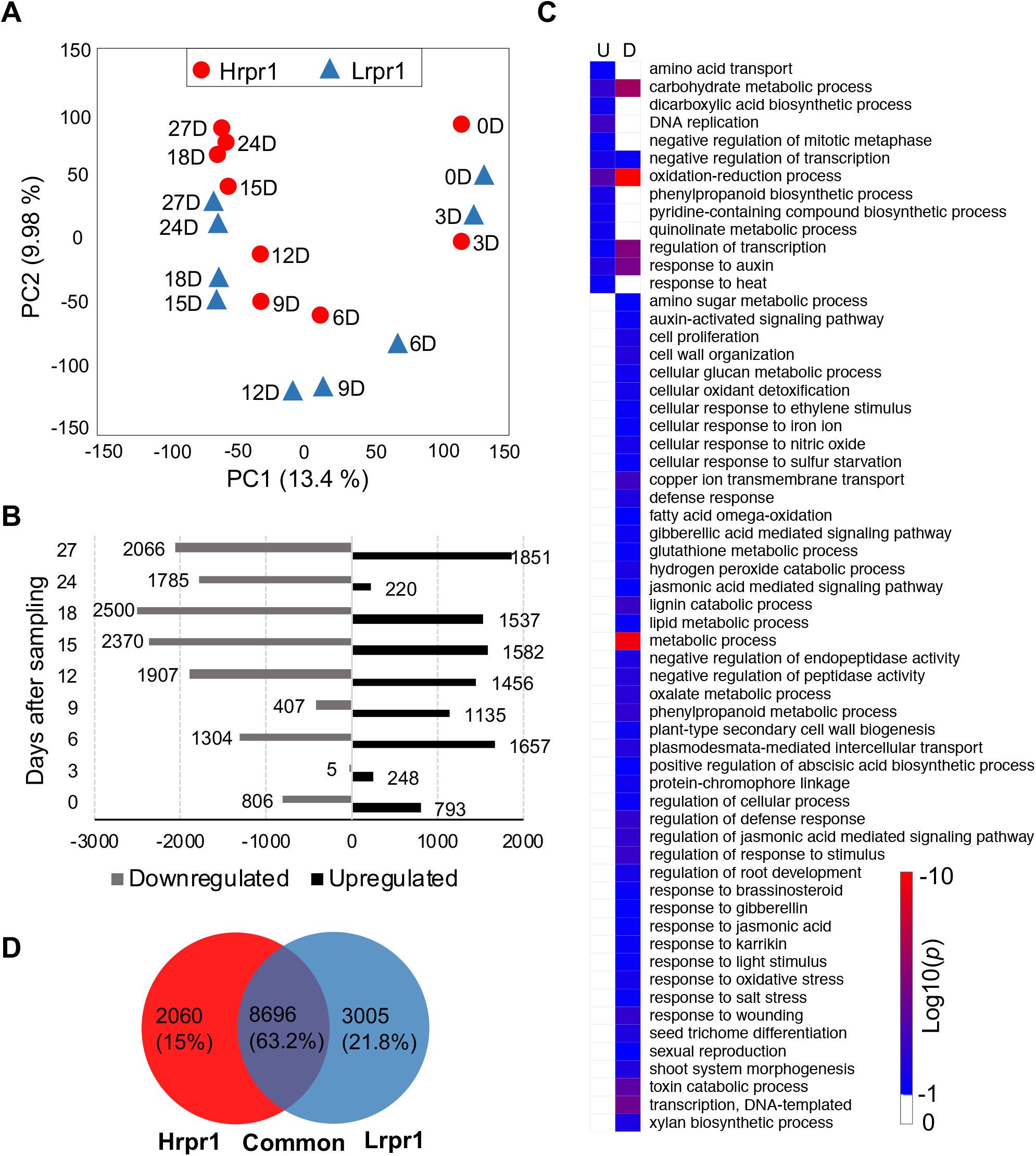
Transcriptomic dynamics during RPR development. (A) Clustering based on PCA performed for different developmental stages of Lrpr1 (blue triangles) and Hrpr1 (red ovals) internodes. The developmental stages (days after sampling; D) are mentioned with corresponding triangles and ovals. (B) Differentially expressed genes during internode development. The *x*-axis represents number of upregulated and downregulated genes in Hrpr1 compared to Lrpr1 and *y-* axis represents developmental stages. (C) GO enrichment analysis showing significantly enriched biological processes with FDR corrected *p* < 0.05, based on downregulated (D) and upregulated (U) genes in Hrpr1 compare to Lrpr1. (D) Unique and common differentially expressed genes in Hrpr1 and Lrpr1.

Comparative analysis of transcriptome changes between Lrpr1 and Hrpr1 at each of the different stages (0 DAS to 27 DAS) showed that 8327 (21%) genes were differentially expressed (DE) during one or more stages (Figure 5B, Table S3). Consistent with phenotypic changes (Figure 3A), a larger number of DE genes were observed after 3 DAS. Furthermore, while the early stages at 3, 6, and 9 DAS were characterized by more DE genes showing upregulation in Hrpr1 relative to Lrpr1 and suggesting higher RPR to be a biologically more active process, this trend was reversed in the later stages. Gene ontology (GO) enrichment analysis of upregulated and downregulated genes identified several important enriched biological processes (GOBP) (Figure 5C). Notable upregulated GOBP in Hrpr1 consisted of phenylpropanoid biosynthesis, amino acid transport, dicarboxylic acid biosynthesis, pyridine-containing compound synthesis, and quinolinate metabolism. Major upregulated GOBP in Lrpr1 consisted of xylan biosynthesis, cellular glucan metabolic process, amino sugar metabolic process, cell wall organization, defense response, gibberellic acid- and jasmonic acid-mediated signaling pathways, lignin catabolic process, secondary cell wall biogenesis, and response to oxidative stress.

To further understand the distinct transcriptome dynamics during the development of the high and the low RPR phenotype in the internodes, we compared the transcriptome at 0 DAS with subsequent developmental stages for each of the inbred lines (Figure S2). The total number of DE genes at various stages of RPR development relative to 0 DAS was similar in Hrpr1 (10758, 27.1%) and Lrpr1 (11701, 29.5%) (Figure S2A-B). However, more DE genes were detected in Hrpr1 relative to Lrpr1 at the early stages of 3, 6, 9 DAS. Furthermore, consistent with the combined analysis (Figure 5B), more DE genes were upregulated that downregulated in Hrpr1 at compared to Lrpr1 at these early stages (Figure S2A-B), further supporting enhanced biological activity underlying high RPR phenotype. Finally, among the DE genes, 2060 (15%) and 3005 (21.8%) were unique for Hrpr1 and Lrpr1, respectively, while 8696 (63.2%) were common for both inbreds (Figure 5D) suggesting that relatively fewer genes are responsible for distinct RPR phenotype. The GO enrichment analysis of unique and common genes showed that oxidation-reduction and phosphatidylcholine metabolic process predominantly enriched in Hrpr1 while nucleic acid phosphodiester bond hydrolysis and RNA modification were enriched in Lrpr1 (Figure S2C).

### Integrating selection mapping with gene expression identified high priority genes associated with RPR

Given the large number of candidate genes obtained from selection mapping and from differential expression in Lrpr1 and Hrpr1, we applied an integrative approach to identify the high priority RPR associated genes. From the SNPs that underwent significant change in frequency during divergent selection, we chose 5% SNPs with the highest *F_ST_* resulting in 183 and 171 high priority SNPs for C15-H and C15-L populations, respectively (Table S4). We mapped these SNPs to nearby two genes within a 100 kb (100 kb on each side) window and further filtered them based on their intersection with differentially expressed genes (fold change >2) between Lrpr1 and Hrpr1 at each of the developmental stages. Remarkably, 98 and 76 genes were found to be associated with low and high RPR, respectively (Table S5). The GO analysis of these genes showed the notable enrichment of pigments and glucuronoxylan synthesis in low RPR phenotype while enrichment of lignin biosynthesis, drought recovery, stress response, and salicylic acid catabolic process in the high RPR phenotype (Figure 6, Table S6).

**Figure 6.**
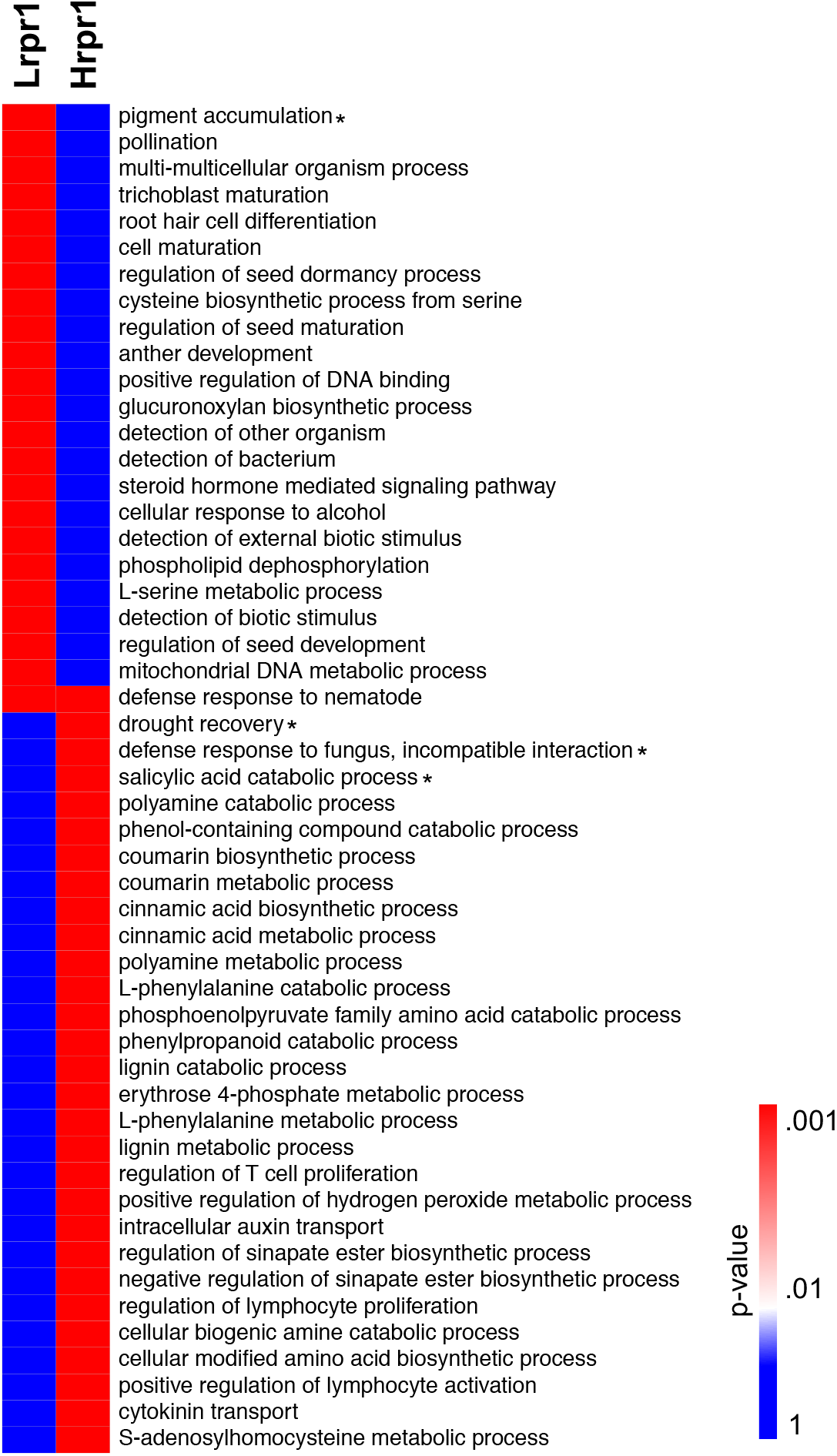
GO terms enriched in Lrpr1 and Hrpr1 based on the genes identified by combining selection mapping and transcriptome analyses. GO terms represented here are significant at *p* < 0.01. Asterisks next to GO term description denote significance after Bonferroni correction.

### Meta-analysis of the genetic architecture of RPR

To generate a comprehensive overview of the genetic architecture of RPR and for the identification of high confidence candidate genes, we combined the findings of our study with other diverse lines of evidence. The criteria used for meta-analysis were 1) location of each of the 354 high priority SNPs (Table S4); 2) genomic regions identified from previous genetic studies including QTL (Flint-Garcia et al., 2003; Hu et al., 2012; Li et al., 2014) and GWAS (Peiffer et al., 2013) analyses; 3) differential expression of the genes identified from overlapping SNPs (Table S3); 4) overlap with genes associated with SNPs enriched in polysaccharide and lignin genome space. Since flanking markers were not available for the QTL derived from MoSCSSS-derived populations (Flint-Garcia et al., 2003), we added 10 Mb windows on both sides of the marker designated as the QTL position in the study. For the NAM GWAS SNPs, genes present within ±100 kb region of associated SNPs were included in this analysis. This analysis identified several key regions, particularly at chromosomes 2, 3, 7, and 9 that overlapped among multiple studies and, therefore, potentially harbor genes and/or regulatory elements associated with RPR (Figure 7). Using at least three of the four criteria defined for meta-analysis, forty candidate genes were identified in these regions (Table 2). For instance, a genomic region on chromosome 2, supported by two additional studies, identified a Myb transcription factor *ZmMYB31* as a candidate gene involved in specifying the high RPR phenotype. Another region on chromosome 3, supported by at least two additional studies, identified a glucosidase encoded by *geb1*. Interestingly, two gene *lic1* and *lic2* identified on chromosome 6 region were supported by an additional study. Another important region on chromosome 9, supported by three additional studies, harbors two candidate genes including a MADSs box gene *ZmMADS1* and an expansin encoded by *Expansin-like A2* (*ZmEXPA2*). Other key candidate genes in the genomic regions identified by at least one more genetic study besides the current study for high RPR include *phenylalanine ammonia lyases* (*ZmPAL7, ZmPAL9*), *Caleosin protein and Xyloglucan galactosyltransferase MUR3*. The important genes identified to be associated with low RPR phenotype and supported by additional genetics studies include *ZmNAC25, receptor-like protein kinase HSL1, ZmWRKY55, DELLA protein RGA (gras46), xyloglucan 6-xylosyltransferase 2* (*ZmGT5*) *histidine kinase5 (hk5), and auxin-responsive protein IAA41 (ZmIAA41)*. The genes identified from the meta-analysis are promising candidates for RPR manipulation in future studies.

**Figure 7.**
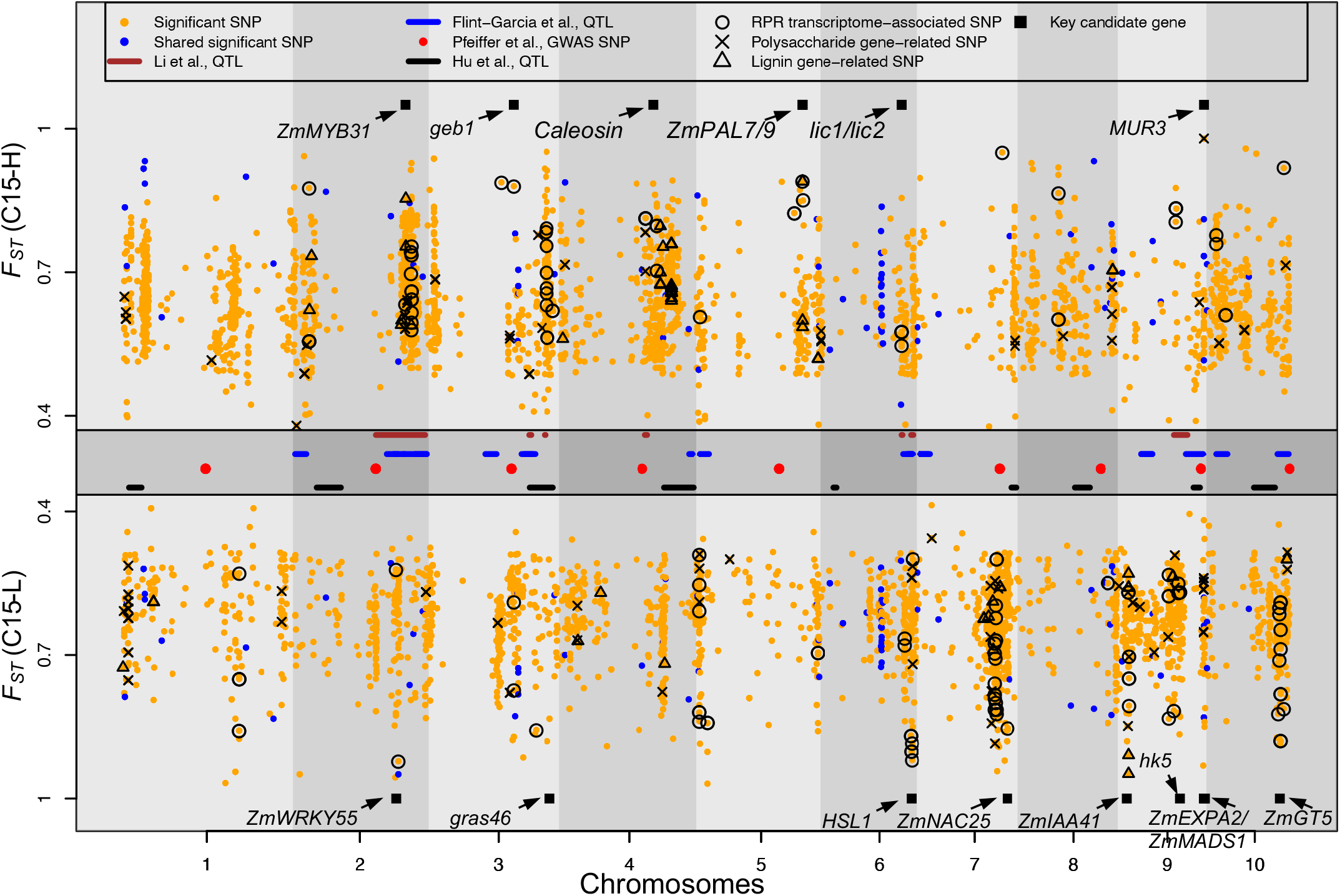
Genetic architecture of RPR deduced from integration of different datasets. Orange dots in the upper and lower panel indicate significantly selected SNPs in high (C15-H) and low (C15-L) RPR, respectively. Middle panel indicates the QTL (blue, brown and black lines) and GWAS SNPs (red dots) associated with RPR reported in earlier genetic studies (Flint-Garcia et al., 2003; Hu et al., 2012; Peiffer et al., 2013; Li et al., 2014). Black circles highlight the SNPs lying in the vicinity of genes with at least 2-fold differential expression in Hrpr1 and Lrpr1 internodes. Crosses, and triangles represent the SNPs associated with polysaccharide and lignin pathways genes, respectively. Key genes associated with RPR identified from different approaches are indicated with black squares.

**Table 2.** Key genes related to rind penetrance resistance identified from the metanalysis. Criteria refer to the four criteria defined in the text that were used to identify candidate genes.

| RPR | SNP position | *F<sub>ST</sub>* | Gene ID | Chr | Gene annotation | Fold change | Criteria |
| --- | --- | --- | --- | --- | --- | --- | --- |
| High | 202,580,532 | 0.63 | Zm00001d006236 | 2 | *ZmMYB31* | 2.6 | 1, 2, 3 |
| High | 197,606,711 | 0.81 | Zm00001d006082 | 2 | *PIN-formed protein11 (pin11)* | 4.9 | 1, 2, 3 |
| High | 210,185,759 | 0.58 | Zm00001d006511 | 2 | *Glucose-6-phosphate/phosphate translocator 2 (gpm334)* | 3.0 | 1, 2, 3 |
| High | 211,751,720 | 0.91 | Zm00001d006591 | 2 | *F-box domain containing protein* | -3.9 | 1, 2, 3 |
| High | 212,878,293 | 0.59 | Zm00001d006627 | 2 | *L-type lectin-domain containing receptor kinase IX.1* | 5.6 | 1, 2, 3 |
| High | 213,548,099 | 0.64 | Zm00001d006653 | 2 | *AP2-EREBP transcription factor (ereb62)* | 4.5 | 1, 2, 3 |
| High | 218,576,041 | 0.67 | Zm00001d006883 | 2 | *Calmodulin binding protein* | -2.6 | 1, 2, 3 |
| Low | 185,589,850 | 0.52 | Zm00001d005741 | 2 | *NAD(P)H-quinone oxidoreductase subunit L* | 6.9 | 1, 2, 3 |
| Low | 186,080,069 | 0.80 | Zm00001d005749 | 2 | *ZmWRKY55* | -4.6 | 1, 2, 3 |
| Low | 189,506,978 | 0.92 | Zm00001d005813 | 2 | *Protein TIFY 10B (zim13)* | 4.5 | 1, 2, 3 |
| High | 153,354,929 | 0.88 | Zm00001d042143 | 3 | *Glucan endo-1,3-beta-glucosidase homolog1 (geb1)* | -4.9 | 1, 2, 3 |
| High | 212,853,636 | 0.82 | Zm00001d043881 | 3 | *Digalactosyldiacylglycerol synthase 1* | 24.2 | 1, 2, 3 |
| High | 213,845,025 | 0.87 | Zm00001d043921 | 3 | *ZmNAC82* | -3.8 | 1, 2, 3 |
| High | 214,419,702 | 0.76 | Zm00001d043947 | 3 | *Lipid phosphate phosphatase 2* | 7.8 | 1, 2, 3 |
| Low | 193,474,769 | 0.86 | Zm00001d043257 | 3 | *G-type lectin S-receptor-protein kinase CES101* | 2.5 | 1, 2, 3 |
| Low | 218,247,550 | 0.83 | Zm00001d044065 | 3 | *DELLA protein RGA (gras46)* | -2.6 | 1, 2, 3 |
| High | 39,730,201 | 0.77 | Zm00001d049678 | 4 | *ZmNAC115* | -5.5 | 1, 2, 3 |
| High | 155,986,150 | 0.81 | Zm00001d051361 | 4 | *VAN3-binding protein* | -4.9 | 1, 2, 3 |
| High | 169,990,594 | 0.52 | Zm00001d051800 | 4 | *Caleosin protein* | -2.5 | 1, 2, 3 |
| High | 191,466,721 | 0.60 | Zm00001d017275 | 5 | *Phenylalanine ammonia lyases (ZmPAL7/ZmPAL9)* | -2.5 | 1, 2, 3, 4 |
| High | 146,400,206 | 0.56 | Zm00001d038049 | 6 | *Lichenase (lic1/lic2)* | -6.4 | 1, 2, 3 |
| Low | 151,942,963 | 0.67 | Zm00001d038221 | 6 | *ZmNAC20* | 4.9 | 1, 2, 3 |
| Low | 164,388,708 | 0.90 | Zm00001d038791 | 6 | *Receptor-like protein kinase HSL1* | 2.4 | 1, 2, 3 |
| Low | 165,372,185 | 0.88 | Zm00001d038831 | 6 | *Type IV inositol polyphosphate 5-phosphatase* | 3.4 | 1, 2, 3 |
| Low | 166,774,503 | 0.89 | Zm00001d038905 | 6 | *putative beta-14-xylosyltransferase IRX10L* | 2.1 | 1, 2, 3 |
| Low | 163,561,967 | 0.85 | Zm00001d021818 | 7 | *ZmNAC25* | -6.9 | 1, 2, 3 |
| High | 103,637,970 | 0.81 | Zm00001d046725 | 9 | *D-mannose binding lectin receptor-like protein kinase* | -3.8 | 1, 2, 3 |
| High | 104,342,643 | 0.83 | Zm00001d046748 | 9 | *Putative polyphenol oxidase family protein* | -6.0 | 1, 2, 3 |
| High | 155,402,586 | 0.66 | Zm00001d048389 | 9 | *Xyloglucan galactosyltransferase MUR3* | 3.6 | 1, 2, 3, 4 |
| Low | 111,179,135 | 0.57 | Zm00001d046928 | 9 | *Histidine kinase5 (hk5)* | -2.4 | 1, 2, 3 |
| Low | 154,806,326 | 0.83 | Zm00001d048344 | 9 | *Expansin-like A2 (ZmEXPA2)* | 2.7 | 1, 2, 3, 4 |
| Low | 156,836,924 | 0.54 | Zm00001d048469 | 9 | *Hydroxylase5 (hyd5)* | -2.9 | 1, 2, 3 |
| Low | 15,993,004 | 0.88 | Zm00001d045203 | 9 | *Aux/IAA-transcription factor 41 (ZmIAA41)* | 25.8 | 1, 2, 3 |
| Low | 156,985,230 | 0.81 | Zm00001d048474 | 9 | *ZmMADS1* | 5.7 | 1, 2, 3 |
| High | 139,066,037 | 0.92 | Zm00001d026127 | 10 | *Patched family protein (mmp71)* | 2.1 | 1, 2, 3 |
| Low | 129,135,133 | 0.82 | Zm00001d025774 | 10 | *Peroxidase superfamily protein* | -2.4 | 1, 2, 3 |
| Low | 130,349,712 | 0.60 | Zm00001d025814 | 10 | *Beta-16-galactosyltransferase (ZmGALT29A)* | -4.0 | 1, 2, 3 |
| Low | 131,515,767 | 0.89 | Zm00001d025851 | 10 | *Xyloglucan 6-xylosyltransferase 2 (ZmGT5)* | -5.6 | 1, 2, 3 |
| Low | 133,094,359 | 0.59 | Zm00001d025886 | 10 | *Cytokinin-O-glucosyltransferase 3* | 8.7 | 1, 2, 3 |
| Low | 138,350,968 | 0.81 | Zm00001d026095 | 10 | *Sugar isomerase (SIS) family protein* | 4.9 | 1, 2, 3 |

## Discussion

Rind strength is an important predictive phenotype of stalk strength and stalk lodging resistance. Rind penetrance resistance (a.k.a. rind puncture resistance) is an effective measure of rind strength that has been shown to be negatively correlated with the stalk lodging incidence in several field studies (Dudley, 1994; Hu et al., 2012; Sekhon et al., 2019). However, lack of knowledge about the genetic mechanisms underlying RPR has been a major obstacle to improving stalk strength and stalk lodging resistance. We employed selection mapping in diverse populations, transcriptomics of inbred lines with extreme RPR phenotypes, metabolite analyses, and combined these findings with the previous genetic studies to provides a comprehensive overview of the genetic architecture of the RPR trait in maize.

### Divergent selection significantly altered allele frequencies at putative RPR-associated regions

Divergently selected populations provide a unique and valuable resource for mining genes and alleles underlying RPR and, therefore, stalk strength. A frequent and valid critique of selection mapping experiments carried out on wild or domesticated populations (Hufford et al., 2012; Beissinger et al., 2016; Plassais et al., 2019) is that these approaches rely on “outlier tests” for identification of divergently selected SNPs/regions. Consequently, these tests are not equivalent to significance tests (Narum and Hess, 2011; Lorenz et al., 2015). To avoid this criticism, we established significance by comparing observed allele frequency differentiation between selected and unselected populations. This approach allowed us to account for differentiation based on genetic drift alone. The rationale for applying this approach is that, like in a subset of other breeding studies carried out within over a short time span of a few to tens of generations on an evolutionary scale (Lorenz et al., 2015), accurate demographic parameters were recorded during the RPR selection experiment (Martin et al., 2004). This approach makes our analysis more comparable to experimental evolution studies in model species (Kofler and Schlötterer, 2014) than to most selection mapping studies in crops (e.g., Zhou et al., 2015; Gage et al., 2018). In contrast to wild populations (Plassais et al., 2019) or domesticated populations studied over an evolutionary timeframe (Hufford et al., 2012; Beissinger et al., 2016), wherein the accurate demographic parameters are unknown, knowledge of such parameters in our study make it straight forward to establish statistical significance via no-selection simulations.

Significant evidence of selection for a large number of SNPs, 3656 for C15-H and 3412 for C15-L, indicates a major change in the genome landscape of the selected population compared to C0 population. Our observations implicating a large number of significant SNPs are consistent with those observed in experimentally evolved populations of model species including drosophila (Turner et al., 2011; Turner and Miller, 2012) and yeast (Araya et al., 2010). Discovery of such a large number of SNPs results, in part, from the fact that the resolution and power of such studies depend heavily on levels of linkage disequilibrium before selection commenced (Kofler and Schlötterer, 2014). Moreover, a SNP with significantly altered allele frequency due to selection is not necessarily causal for the trait of interest, and may instead be linked to a causal site through a phenomenon known as hitchhiking (Smith and Haigh, 1974).

Although our analysis showed significant enrichment for SNPs selected in the high and low RPR directions, the number of overlapping SNPs (110) was a small fraction of the total number of significant SNPs. There are several possible reasons, both practical and biological, for this outcome. First, selection mapping evaluated by quantification of the change in allele frequency, as implemented here, is most powerful to identify SNPs that have moved from a low frequency to a high frequency due to selection, or vice versa (Vlachos and Kofler, 2019). Therefore, even when selection operated on the same SNP in both populations, our study may only have the power to detect the SNP the population where the frequency of the SNP was significantly altered from C0. Second, our drift simulation approach for declaring significance included only single SNP tests and, therefore, it is possible that different SNPs associated with high and low RPR in the same genomic region. Indeed, manual inspection of significantly selected SNPs in either direction identifies several hotspots of SNPs in the high and low RPR that may correspond to selection on the same underlying gene (Figure 7). Third, the relatively small amount of overlap may indicate that distinct biological processes underlie high and low RPR. This argument is supported by the enrichment of significant SNPs in the lignin genome space in the high RPR direction and enrichment in the polysaccharide genome space in the low RPR direction.

### Transcriptomic analysis reveals both expected and novel process related to RPR

Consistent with large changes in the genome landscape indicated by selection mapping, high levels of biological activity associated with RPR is evident from differential expression of 21% of the annotated maize genes. Global transcriptome analysis showed that the high and low RPR phenotypes start to diverge very early during the development of an internode. While differentiation of secondary cell wall has been considered as the major contributor to strength, such differentiation happens after the internode elongation is complete (Cosgrove and Jarvis, 2012; Zhang et al., 2019). Therefore, high transcriptome activity at early stages indicates that differences in RPR are not limited to secondary cell wall biosynthesis. Furthermore, over 2000 and 3000 unique DE genes associated with high and low RPR, respectively, outnumber 845 genes associated with primary and secondary cell wall synthesis. While this gap will shrink with the discovery of additional genes related to cell wall synthesis, and cell wall is one of the most important determinants of RPR, identification of novel biological activities and processes will further elucidate the determination of rind strength.

GO enrichment based on DE genes provides a global view of the key processes and activities involved in the determination of rind strength. In the internodes of high RPR inbred, Hrpr1, synthesis of secondary metabolites is a prominent process as indicated by enrichment of phenylpropanoid biosynthetic processes. Phenylpropanoid pathway is responsible for a large number of natural plant products including lignin, coumarins, flavonoids, anthocyanins, phenolic acids, and stilbenes (Vogt, 2010). Besides providing resistance to various biotic and abiotic stresses, these compounds are needed for reinforcing the cell wall and, therefore, providing mechanical strength to plants (Dixon et al., 2002; Vogt, 2010; Fraser and Chapple, 2011; Gray et al., 2012). Enrichment of quinolinate metabolism indicates increased synthesis of NAD, which is known to facilitate lignification and oxidative cross linking of lignin and polysaccharides in the cell wall (Pétriacq et al., 2012; Higuchi et al., 2014). Enrichment of amino acid transport suggests an important albeit less understood role of this process in the determination of RPR. Amino acids are involved in the synthesis of cell wall protein and enzymes, and act as the precursors for monolignol synthesis (Maeda and Dudareva, 2012).

In the transcriptome of low RPR inbred, Lrpr1, enrichment of xylan and glucan metabolism supports higher flux of photoassimilates towards polysaccharides which, together with lower lignification, explains higher recovery of cell wall polysaccharides. Enriched metabolism of amino sugars indicates a higher accumulation of glycoproteins that may be important for providing mechanical strength in the absence of optimal lignification. Amino sugars are important components of chitin and recent studies suggest their role in the determination of plant architecture (Vanholme et al., 2014). Enrichment of auxin signaling, and response to ethylene, brassinosteroids (BR), jasmonic acid, and gibberellic acid in Lrpr1 internodes indicates important, yet sparsely understood, role of hormones in the determination of RPR. In Arabidopsis, BR are known to regulate cellulose synthesis, and defects in BR synthesis and signaling result in impaired cellulose accumulation in the cell walls (Xie et al., 2011; Sánchez-Rodríguez et al., 2017). Auxin concentration is linked to cell expansion during cell wall synthesis and thereby affects the mechano-chemical aspects of plant development (Braybrook and Peaucelle, 2013; Paque et al., 2014; Lehman et al., 2017; Majda and Robert, 2018). Gibberellic acid has also been found to regulate cell wall extension and, interestingly, enhance the lignification of xylem (Bai et al., 2012; Falcioni et al., 2018).

### Selection on high and low RPR operated on distinct gene sets

Selection mapping, transcriptomic, and metabolic analyses indicate that plants with high and low RPR employ distinct approaches to partitioning photoassimilates towards cell wall biosynthesis. The annotation of the cell wall genes and examination of their enrichment during divergent selection revealed that selection preferentially acted on polysaccharide synthesis genes during selection for low RPR and lignin biosynthetic genes during selection for high RPR. This altered genome landscape is well reflected in the transcriptional activity and chemical composition of the internode cell walls. The enrichment of the lignin pathway during selection for high RPR phenotype is consistent with elevated expression of lignin biosynthetic genes and the higher lignin content in Hrpr1. The enrichment of polysaccharides pathway during selection for low RPR is consistent with lower lignification and an increased amount of extractable sugars from the cell wall in Lrpr1.

Chemical composition of rind cell walls, which generally refers to lignin and polysaccharides, is an important determinant of stalk strength (Zuber et al., 1980; Appenzeller et al., 2004). However, whether these constituents impart a positive or negative impact on stalk strength is debated (Sekhon et al., 2019). Cellulose has been proposed to either positively associate with stalk strength or to be no consequence (Appenzeller et al., 2004; Ye et al., 2016). Reduction in lignin content, observed in the maize brown midrib (bmr) mutants, has been associated with decreased stalk strength (Miller et al., 1983). Study of a monocot-specific microRNA (ZmmiR528) also showed that increased lignification is positively associated with improved RPR (Sun et al., 2018). The current study clearly indicates the opposing effect of the accumulation of lignin and polysaccharides on RPR. Attempt to relate the gross amount of polysaccharides and lignin with RPR or stalk strength, however, is an over-simplification of a complex structural phenomenon. The microstructure of these components and the effect of interactions between these constituents must be considered to understand their effect on stalk strength.

While considering the lignification of secondary cell walls, the emphasis is often placed on the content of the three major monolignols, and the processes involved in the polymerization of these lignin monomers is overlooked (Tobimatsu and Schuetz, 2019). Detailed metabolic analysis of Hrpr1 and Lrpr1 internodes highlights the role of additional phenolics in cell wall lignification especially during the early stages of internode development. Tricin, a monolignol that provides nucleation sites for incorporation of the three major monolignols and initiates lignification of grass cell walls (Lan et al., 2015), emerged as an important determinant of lignification associated with RPR. High tricin content would result in more lignin chains, and interaction of these chains with each other and with other molecules (e.g. hemicellulose) would result in higher rind strength. Consistent with this notion, higher FA and pCA content associated with high RPR further indicates that higher cross-linking, and polymerization of lignin and other constituents is important for high rind strength.

### A roadmap to enhanced mechanistic understanding and exploitation of RPR for improving lodging resistance

While ‘stand-alone’ omics investigations, including those looking at genome or transcriptome landscapes underlying a complex trait, provide useful information about the underlying biology and genetics, results from such studies are also often riddled with false positives and false negatives. Integrating selection mapping and transcriptome analyses with previously reported QTL and GWAS studies provides a more compressive view of the genetic architecture of RPR and allows the identification of a core set of candidate genes (Figure 7). Several genomic hotspots detected by examining selection signatures in the divergently selected RPR populations that co-localize with previously reported regions likely harbor large effect genes. Conversely, regions exclusively detected by selection mapping capture the allelic variation present in the inbred lines used to develop the base (C0) population.

Many of the promising candidate genes highlighted by the integrated analysis are associated with cell wall biology. For instance, both selection mapping and transcriptome analyses identified a region on chromosome harboring two genes (*ZmPAL7, ZmPAL9*) encoding for phenylalanine ammonia lyase which catalyzes the first reaction of the phenylpropanoid pathway and regulates the flux towards the synthesis of all secondary metabolites (Vanholme et al., 2010). Impairment of phenylpropanoid biosynthesis in *pal1* and *pal2* mutants, leads to a reduced amount of lignin and decreased cell wall strength in Arabidopsis (Rohde et al., 2004). Low RPR phenotype and low lignin content are associated with reduced expression of *ZmPAL7* and *ZmPAL8* (Sun et al., 2018). Likewise, a candidate transcription factor, ZmMYB31, on chromosome 2 has been established as a regulator of lignin biosynthesis in maize (Fornalé et al., 2006). Overexpression of *ZmMYB31* in Arabidopsis reduced the lignin biosynthesis but did not alter the composition of the lignin polymer (Fornalé et al., 2010). Of multiple candidate genes present on chromosome 9, a MADS-box transcription factor encoded by *ZmMADS1* has been implicated in auxin transport and signaling (Khanday et al., 2013) further supports the role of auxin in the determination of RPR. Putative orthologs of two other candidate genes, *ZmNAC25* and *ZmEXPA2*, have been shown to be involved in cell expansion in Arabidopsis through GA/DELLA-NAC25/NAC1L-EXPA2 regulatory network (Sánchez-Montesino et al., 2019). Given that GA signaling has multiple regulatory effects on cellulose synthesis (Huang et al., 2015; Felipo-Benavent et al., 2018; Sánchez-Montesino et al., 2019), the role of *ZmNAC25* in RPR warrants detailed investigations. Finally, while some of the genes identified in this study can be directly or indirectly linked with some aspects of stalk strength, many lacks such association due to absence of experimental evidence. These genes have the potential to provide a wealth of information related to the biology of stalk strength in maize and related grasses.

## Conclusions

Despite technological advances, progress in deciphering the genetic architecture of complex traits through omics approaches has been slow. In the case of stalk lodging resistance, a trait impacted by a number of internal and external factors, lack of reliable and scalable methods for phenotypic evaluation of the germplasm adds additional challenges. To this end, systematic analysis of populations divergently selected for RPR, a reliable predictor of lodging resistance, offers a unique approach to generate novel insights into the molecular genetic mechanisms governing stalk strength. Combining the results from selection mapping with those from the current state-of-the-art approaches provide higher quality candidate mechanism and genes that will guide functional studies to understand the molecular architecture of RPR and stalk strength. Furthermore, the valuable information generated by this study will boost efforts for genetic improvement of stalk strength in maize, sorghum, and other grasses.

## Materials And Methods

### Genetics stocks

The MoSCSSS C0 population, hereafter called C0, was derived from 14 inbred lines (A657, A632Ht, B14AHt, B37Ht, B68, B73, B76, B84, CM105, H84, N28Ht, N104, Oh514, and Pa864P) that were either direct derivatives or closely related to the Iowa Sty Stalk Synthetic heterotic group (Martin et al., 2004). Populations derived from 15 cycles selection for high and low RPR by Dr. Larry Darrah and colleagues at the University of Missouri (Figure 1A) along with the C0 population were kindly provided by Dr. Sherry Flint-Garcia. The high and low populations, originally named as MoSCSSS H24 (High Rind Penetrometer [HRP]) C15 and MoSCSSS H25 (Low Rind Penetrometer [LRP]) C15 were renamed as C15-H and C15-L, respectively (Figure 1). Each of these three populations was multiplied separately by making 250 unique crosses within the population such that each of the 500 parent plants was either used as a male or female only once, and by bulking 100 seeds from each of the resulting crossed ears.

For the development of inbred lines, one randomly chosen plant each from C15-H and C15-L populations was self-pollinated. A single row of 15-20 plants was grown from each resulting ear, and one randomly chosen plant from each row was again self-pollinated. This process was repeated six times to attain near-homozygosity. Seed from each ear was then used to multiply seed for each of the inbred lines. The inbred lines derived from C15-H and C15-L populations were designated as Hrpr1 and Lrpr1, respectively.

### Measurement of rind penetrometer resistance

For phenotypic analysis, the Hrpr1 and Lrpr1 inbred lines were grown in a randomized complete block design with three replications at Clemson University Calhoun Field Laboratory, Clemson, SC in summer 2017. In each block, a four-row plot of each inbred line was grown with row length and row-to-row distance being 4.57 meters and 0.762 meters, respectively, resulting in a total pot size of 13.93 m^2^. Data was recorded on 12th internode as, on an average, this internode lies below the primary ear-bearing node in these inbred lines. As each maize leaf is attached to a node, leaves were used to identity the 12^th^ internode. Since the juvenile leaves senesce and become undetectable at later developmental stages, at the vegetative 4 (V4) stage marked by the presence of four fully extended leaves, a hole was punched in the fifth leaf and this leaf was subsequently used to identify the 12^th^ internode. The development of the 12^th^ internode was visually followed by dissecting plants every 3 days. The phenotypic data and sample collection were started at the V8 stage attained 45 days after sowing (DAS) when the 12^th^ internode was approximately 0.5 cm long. The samples were collected at a three-day interval. RPR was measured with a 2 mm diameter probe with flat tip (49FL81, Grainger Inc., Lake Forest, IL) installed on an Imada^®^ Digitial Force Gauge (ZTA-DPU). A steel plate covering the base of the probe was installed on the force gauge to prevent secondary contact after the initial puncture. For the C0, C15-H, and C15-L populations, RPR was recorded on the internode below the primary ear-bearing node after careful removal of the leaf sheath. The puncture test was performed in the center of the internode, perpendicular to the minor axis of the stalk cross-section (i.e., in the direction of the major axis of the stalk cross-section).

### Genotyping-by-sequencing and SNP calling

Individual leaf samples from 96 plants of each of the three populations (C0, C15-L, and C15-H) were collected from field-grown plants at the V8 stage, flash-frozen in liquid nitrogen, and stored at −80°C. DNA of each plant was extracted using the cetyl(trimethyl)ammonium bromide (CTAB) method (Saghai-Maroof et al., 1984). DNA was cleaned using a modification of the Qiagen DNeasy 96 Plant kit (Qiagen, Germantown, MD), diluted to 20ng/μl, and approximately 200ng DNA was used for modified *ApeKI* genotyping-by-sequencing (GBS) (Elshire et al., 2011). The modification included separating each 96 well plate into 4 pools of 24 adapter-ligated samples. Barcodes were designed on DeenaBIO and synthesized by IDT (Coralville, Iowa, USA. Pre-polymerase chain reaction of the pooled sample and PCR amplification was done using ThermoFisher Phusion II master mix. Quantification of each enriched library pool was done by Qubit (Invitrogen, Carlsbad, CA) and distribution was analyzed on Agilent Bioanalyzer (Agilent Technologies, Santa Clara, CA) high sensitivity DNA chip. All 288 individually barcoded samples were pooled. NextSeq High Output single-end 75-bp sequencing was performed at University of Missouri, Columbia DNA Core (https://dnacore.missouri.edu/).

### SNP calling, filtration, and estimating allele frequency

Sequenced reads were aligned to the maize B73 reference genome version 4 (AGPv4) (Jiao et al., 2017) using the Tassel 5 GBS v2 pipeline (Glaubitz et al., 2014). This alignment resulted in a total of 219,758 initial SNP markers with the mean read depth of ~2.6x. These SNPs underwent very stringent filtering due to the fact that GBS often yields a high proportion of missing data (Beissinger et al., 2013). First, only SNPs with two alleles at a particular locus were included. Individuals were then filtered to remove SNPs with greater than 40% of markers missing, resulting in the removal of 37 individuals from C0, 15 individuals from C15-H, and 31 individuals from C15-L population. Furthermore, SNPs missing at greater than 50% of observations were also filtered from each population. The resulting SNPs were further filtered to remove low confidence SNPs with read count less than 10. After filtering, 159,849 SNPs were retained from 205 individuals for the downstream analysis. Reference allele frequency was computed in each population using VCFtools (Danecek et al., 2011).

### Detection of selected loci

Two techniques were used to identify putatively selected regions of the genome. We used a pairwise estimate of *F_ST_* calculated in R version 3.4.1 (R Core Team, 2013) based on a published method (Weir and Cockerham, 1984) and the published analysis pipeline (Beissinger et al., 2014), such that

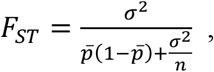

where σ^2^ is the sample variance between the populations, 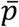 is the average allele frequency between two populations, and *n* is the number of populations. For every locus, a single *F_ST_* value was computed.

*F_ST_* depends on initial allele frequency (Jakobsson, Edge, & Rosenberg, 2013) and cannot alone be used to generate significance thresholds (Akey, 2009). Therefore we implemented a simulation-based test that incorporated the demographic history of the C15-H and C15-L populations to allow for significance testing. Our strategy involved comparing the observed allele frequency divergence at each site to the expected magnitude of genetic drift over the course of the experiment. We simulated a drift-only breeding program based on selection for high and low RPR and evaluated the distribution of allele frequencies after breeding. In our simulations, breeding proceeded for 15 generations with 60 males and 120 females mated each generation in agreement with the scheme used for the development of these populations (Martin et al., 2004). Furthermore, our simulation strategy incorporated sampling of 96 individuals from generation 0 (G0) and G15 for genotyping. To implement these simulations, first we randomly sampled 96 individuals from a pool with allele frequency set between 0.01 and 0.99 with increments of 0.01. This sampling was conducted 1 million times for each initial frequency. Next, the frequency of each random sample was used as an initial frequency for 15 generations of drift simulations, with 60 contributing males and 120 contributing females each generation. The frequencies at the end of G15 were then used for a final sampling of 96 individuals. We used the final simulated frequencies to calculate the 1e-7 and 1-1e-7 quantiles of allele frequency change. Then, we compared the observed pre-selection allele frequencies in C0 to the postselection allele frequencies in C15-L and C15-H to identify SNPs that changed in frequency more than that expected under drift and sampling alone in each population. SNPs with minor allele frequency below 0.01 in C0 were not included in this analysis resulting in 158,196 SNPs. The SNPs with allele frequency change greater than the 1e-7 and 1-1e-17 quantiles are putative candidates for selection, with a 1e-7 false positive rate. Since we evaluated 158,196 SNPs, the experiment-wide false positive rate was fewer than one SNP. Further details, including our simulation scripts, are provided as Supplemental Scripts.

### Annotation of cell wall genes

The cell wall related protein sequences involved in the biosynthesis of cellulose, hemicellulose, and lignin reported in an earlier study (Guillaumie et al., 2007) were used as query in BlastP (evalue 10^-5^) against the latest maize ensemble protein database (version B73_RefGen_v4) to identify complete gene families. The top hit sequences were confirmed and annotated by searching against the Conserved Domains Database (www.ncbi.nlm.nih.gov/Structure/cdd/wrpsb.cgi) and NCBI non-redundant database using evalue of 10^-5·^ The confirmed cell wall genes were analyzed for their associated SNPs enrichment during divergent selection for low and high RPR.

### Testing for the enrichment of selected SNPs within lignin and polysaccharide synthesis genes

A resampling analysis was conducted to test the hypothesis that subsets of genes related to the lignin and the polysaccharide biosynthetic pathways experienced elevated levels of selection as compared to the rest of the genome. For this test, the number of significant SNPs within the total genome space of annotated genes in lignin and polysaccharide biosynthetic pathways were counted. The genome space was defined as positions from 5 kb upstream of the AGPv4 predicted transcription start position to 5 kb downstream of the predicted transcription stop position of the lignin and polysaccharide biosynthetic pathways. All SNPs within these spaces were designated as potentially related to lignin or polysaccharide synthesis, respectively, and subjected to enrichment analysis. Separate counts of the number of lignin and polysaccharide synthesis associated SNPs were calculated for selection in the high and low RPR directions. Then, 1,000 permutations were conducted, where for each permutation, SNP *p*-values (calculated based on allele frequency change) were shuffled over all SNPs positions to generate a set of *permuted-significant* SNPs. For each permutation, the number of *permuted-significant* SNPs within the genome space corresponding to the lignin and the polysaccharide pathway was counted. Based on these permutations, we computed 95% and 99% confidence intervals for the number of significant SNPs that should fall within each genome space if it was not enriched for significant SNPs. We compared the permutation distribution to the observed number of significant SNPs in the lignin and polysaccharide pathway genome space to identify one-sided *p*-values for enrichment.

### Histochemical staining

Mature maize stems were hand sectioned with a razor blade and the cross-sections were stained in phloglucinol-HCl satin following a published protocol (Ruzin, 1999). Samples were imaged with a Zeiss Axioplan2 imaging microscope.

### Tissue collection for metabolic analysis and RNA sequencing

The 12^th^ internode used for measurement of RPR was used for metabolic analysis. All samplings were performed between 8:00 AM and 12:00 PM. At each stage and for each inbred line, three biological replicates of each tissue were collected. A biological replicate consisted of pooled tissue from two different, competitive (non-border and well-spaced from neighbors) plants harvested from the same block and dissected to separate the 12^th^ internode. Samples were chopped into small pieces, flash-frozen in liquid nitrogen, and stored at −80°C.

### Quantification of polysaccharides

For metabolic analyses, the samples were ground to very fine powder under liquid nitrogen with a cryogenic grinder (6875 Freezer/Mill, Spex SamplePrep, Metuchen, NJ) using a 45 second pulse followed by 30 seconds resting period and another 45 second pulse. The resulting fine ground powder was transferred to 15 ml tubes, freeze-dried, and used for various metabolic analyses. The glucose and pentose monomer in cell wall were measured according to a published protocol (Santoro et al., 2010) with modifications described previously (Sekhon et al., 2016).

### Phenolics/lignin extraction

Sequential extraction was used for the separation of phenolic compounds according to published protocol (Li et al., 2015) with modifications. The tricin was extracted with methanol, the ester/ether-bound phenolics were recovered following a mild-base hydrolysis at high temperature, and monolignols were extracted after oxidative degradation of lignins at a higher temperature and pressure in the presence of CuO (Hedges and Ertel, 1982). For quantifying the extractable phenols, 500 mg of freeze-dried tissues was extracted with 2.5 ml of methanol for 3 h with shaking at room temperature. After centrifugation at 1500 *g* for 15 min, 1.5 ml of supernatant was transferred to a glass vial and stored at −20°C for tricin quantification. The remaining pellet was washed twice with 2.5 ml of methanol, dried overnight and, hydrolyzed with 6 ml of 1M NaOH in a glass tube, sparged with Ar, and incubated at 90°C for 3 h in the dark. After cooling on ice, the supernatant was used for the analysis of ester/ether-bound phenolics. The pH of the supernatant was reduced to < 2 using 50% HCl, and the resulting precipitate was discarded after centrifugation. The solution in the tubes was extracted with 2 ml of ethyl acetate and ethyl acetate layer was transferred into GC vials and stored at −20°C for further analysis.The residual pellet was washed twice with 5 ml of deionized water, dried, and stored at −20°C for lignin analysis.

The lignins were depolymerized in 23 ml Acid Digestion Vessels (model 4749 Parr, Instrument Co., Moline, IL, USA) (Kaiser and Benner, 2012). 500 mg of CuO, 75 mg Fe(NH_4_)_2_(SO4)_2_·6H_2_O and 5 ml of freshly prepared 2 M NaOH (pre-sparged with Ar for 30 min) was combined with the base hydrolyzed pellet in Teflon cups. The vessel was sealed and incubated at 155°C for 160 min and then immediately cooled in an ice bath to room temperature. The digestion mixture was transferred to fresh tubes and 50 μl of ethyl vanillin (400 mg l^-1^) was added as an internal standard. The pH of the solution was reduced to < 2 by adding of H2SO4 and mixed gently. After centrifugation, 8 ml of the supernatant was transferred to a clean glass tube, and the lignin-derived phenols were extracted by liquid-liquid partitioning with 2 ml of ethyl acetate.

### Tricin analysis using LC-MS/MS

The methanol extracts were profiled using a quadrupole-orbitrap-iontrap mass spectrometer (Orbitrap Fusion™ Tribrid™ Mass Spectrometer; Thermo Fisher Scientific, Waltham, MA, USA). Compounds were separated on a C18 reversed-phase column (ACQUITY HSS T3, 150 x 2.1 mm, 2.6 μm) using an UHPLC (Dionex Ultimate 3000), with gradient elution employing binary solvent system involving 0.1% formic acid and acetonitrile. The gradient of acetonitrile mix was increased from the initial proportion of 5% to 95% over 25 min and held for 5 minutes before re-equilibrating the column to the initial solvent condition. The solvent flow rate was maintained at 0.15 mL/min, and the column was maintained at 35°C. The samples were introduced into the mass spectrometer through a heated electrospray ionization (H-ESI) interface operated in a negative ionization mode. The Orbitrap Fusion mass spectrometer was operated in high-resolution (120,000 FWHM at 200 m/z) full-scan mode (120-1000 m/z). The optimized parameters were set as follows: sheath gas 60, auxiliary gas 15, sweep gas 1, spray voltage 3.5 kV, probe temperature 350°C, and transfer capillary temperature 300°C. The fragmentation was achieved via collision-induced dissociation with collision energy set at 35%. From the full spectral scans, the extracted ion chromatograms of tricin were derived using theoretical exact masses of [M-H]-ion with 4 significant figures and a scan width of ± 1.0 ppm mass error.

### LC-MS Quantitation of Lignin Phenols

The phenolic monomers resulting from the base hydrolysis of bound phenolics and CuO oxidation of lignins were measured using liquid chromatography coupled to a triple quadrupole mass spectrometer (LC-MS/MS Shimadzu LC-20AT + Shimadzu 8040 mass analyzer; Kyoto, Japan). Ethyl acetate partitioned samples were dried under nitrogen gas stream and re-dissolved in methanol (300 μl volume maintained) for LC-MS/MS analysis. Compounds were separated on a C18 reversed-phase column (Kinetex XB-C18, 150 x 3 mm, 2.6 μm) using a binary solvent gradient. Solvent A was water containing 0.1% formic acid and solvent B was acetonitrile. The solvent flow rate was maintained at 0.3 mL/min and the gradient of acetonitrile was increased from the initial proportion of 25% to 95% over eight minutes on a linear gradient and held for one minute before re-equilibrating the column to initial solvent condition. The samples were introduced into the mass spectrometer through an electrospray ionization (ESI) interface operated in a negative ionization mode. The ESI interface parameters were set as follows: nitrogen nebulizing gas 3 L/min, nitrogen drying gas 12 L/min, capillary voltage −3.5 kV, heat block 400°C, and desolvation line temperature 250°C. The triple quadrupole mass spectrometer was operated in MS/MS mode based on the following multiple reaction monitoring parameters that were optimized using authentic standards: p-coumaric acid (162.8 > 119.1, 92.95, 117.1), dihydroxybenzoic acid (152.9 > 109.05, 65, 67.05), ferulic acid (192.9 > 134.1, 178.1, 149.2), p-hydroxybenzoic acid (136.9 > 93.1, 64.95, 41.1), p-hydroxybenzaldehyde (121 > 92.05, 93, 41.05), p-hydroxyacetophenone (134.95 > 92.05, 93, 120.05), syringic acid (196.85 > 182.1, 123.1 95), syringaldehyde (180.85 > 166.1, 151.1, 123), acetosyringone (194.9 > 180.2, 165.1, 137), vanillic acid (166.85 > 152.05, 108.05, 123.1), vanillin (150.95 > 136.1, 92.05, 108.05), and acetovanillone (164.95 > 150.05, 122.05, 79). Quantitation was performed using a series of 13-point external calibration curve ranging from 12 ppb to 12.5 ppm containing a mixture of all 12 compounds.

### RNA Sequencing

RNA was extracted from the frozen chopped internode samples using TRIzol reagent (Invitrogen, Carlsbad, CA, USA) following the manufacturer’s protocol. The mRNA isolation, cDNA synthesis, and library preparation were performed by Novogene Corporation (Chula Vista, CA) following the standard Illumina (Illumina Inc., San Diego, CA) protocol. The 150 bp paired-end reads generated after sequencing by Illumina HiSeq platform were trimmed of low-quality bases and adapter sequences with the trimmomatic (Bolger et al., 2014). Reads were aligned to reference maize genome (Zea_mays.AGPv4.40.gtf) (Jiao et al., 2017) with the Tophat2 v. 2.1.1 (Trapnell et al., 2012) using Bowtie2 (v. 2.3.4.1) (Langmead and Salzberg, 2012). Cufflinks v.2.2.1 was used to determine Fragments per kilobase per million (FPKM) (Trapnell et al., 2012). The read count was determined with the HTSeq-count v.0.10.0 (Anders et al., 2015). The differential expression between different pairs of stages and genotypes (Hrpr1 and Lrpr1) were calculated using DESeq2 (Love et al., 2014). Genes with *p* ≤ 0.05 were corrected by the false discovery rate (FDR) < 0.05 (Benjamini and Hochberg, 1995). Genes with log2 fold change > 2 and FDR corrected *p* < 0.05 were selected for subsequent analyses. Gene ontology (GO) enrichment was carried out with the GOSeq software tool (Young et al., 2010) and enriched categories were filtered by FDR corrected *p* < 0.05.

### Integrating of selection mapping high priority SNPs and gene expression

The top 5% *F_ST_* SNPs for high and low RPR were mapped to nearby genes using the SNP-to-gene mapping tool implemented in Camoco (v0.6.3) (Schaefer et al., 2018), using the parameters of selecting up to 2 flanking genes within 100 kb windows both upstream and downstream of SNPs. Candidate genes were filtered to only genes that were statistically significant and at least 2-fold differentially expressed between Hrpr1 and Lrpr1 internodes. Gene ontology (GO) enrichment was performed using the hypergeometric statistic from utilities within Camoco to identify the putative biological function of DE genes near extreme *F_ST_* SNPs.

### Analysis scripts

All scripts used for selection signature analysis are available as supplemental files. Drift simulations were performed using scripts Supplemental Script 1 and Supplemental Script 2 that were extracted from beissingerlab.github.io and originally published elsewhere (Lorenz et al., 2015). Significance thresholds from drift simulations were compiled using Supplemental Script 3. Supplemental Script 4 was used for SNP calling and allele frequency calculations. *F_ST_* was calculated using Supplemental Script 5 that was extracted from beissingerlab.github.io/Software/ and originally published elsewhere (Beissinger et al., 2014). Supplemental Script 6 was used to compare changes in allele frequency to drift simulations to determine significance, to conduct bootstrap analysis of the number of overlapping SNPs selected by chance, to perform enrichment analysis testing for selection on polysaccharide and lignin genes, and to visualize the metaanalysis (Figure 7).

### Accession Numbers

All sequence reads are available at Sequence data from this article can be found at the National Center for Biotechnology Information Sequence Read Archive (https://www.ncbi.nlm.nih.gov/sra) under accession number PRJNA623937 (https://dataview.ncbi.nlm.nih.gov/object/PRJNA623937?reviewer=su7vgercui58lfpvquk278b13t)

## Acknowledgments

Authors thank Dr. Sherry Flint Garcia for providing the divergently selected RPR populations, Kate Guill for genotyping, Jackyung Yi, Arlyn Ackerman, and a number of talented undergraduate students for help in the collection of data and samples. Funding for this study was provided by the National Science Foundation grant #1826715 to R.S.S. Support for SNP genotyping included resources from the USDA-ARS, Plant Genetics Research Unit.

## Supplementary Data

**Table S1**. Significant SNPs divergently selected for high RPR (C15-H) and low RPR (C15-L) relative to C0

**Table S2**. Annotation of maize cell wall synthesis genes

**Table S3**. Differentially expressed genes (fold change >2; *p*(adj) < 0.05) in different development stages (DAS) of Lrpr1 compared to Hrpr1

**Table S4**. High priority SNPs selected based on 5% *F_ST_* from divergent selection for high RPR (C15-H) and low RPR (C15-L)

**Table S5**. Genes associated with RPR identified from integration of selection mapping and transcriptome data

**Table S6**. Gene ontology (GO) enrichment of genes identified by integration of selection mapping and differential gene expression

**Figure S1**. Individual monolignols constituents in internodes of Lrpr1 and Hrpr1 inbreds. Shown here are syringic acid, syringaldehyde and acetosyringone of S lignin (A-C); and vanillic acid, vanillin and acetovanillone of V lignin (D-F); p-hydroxy benzoic acid, p-hydroxy benzaldehyde, and p-hydroxy acetophenone of H lignin (G-I).

**Figure S2**. Transcriptomic changes during Lrpr1 and Hrpr1 development. Differentially expressed (DE) genes at each developmental stage relative to 0 DAS in Hrpr1 (A) and Lrpr1 (B). The *x*-axis represents number of upregulated and downregulated genes relative to 0 DAS and *y*-axis represents different stages. (C) GO enrichment analysis showing significantly enriched biological processes (FDR corrected *p* < 0.05) based on unique DE genes in Hrpr1 and Lrpr1, and common DE genes in both inbred lines.

