## Supplementary material for "Genetic Architecture of Maize Rind Strength Revealed by the Analysis of Divergently Selected Populations": Figure S1

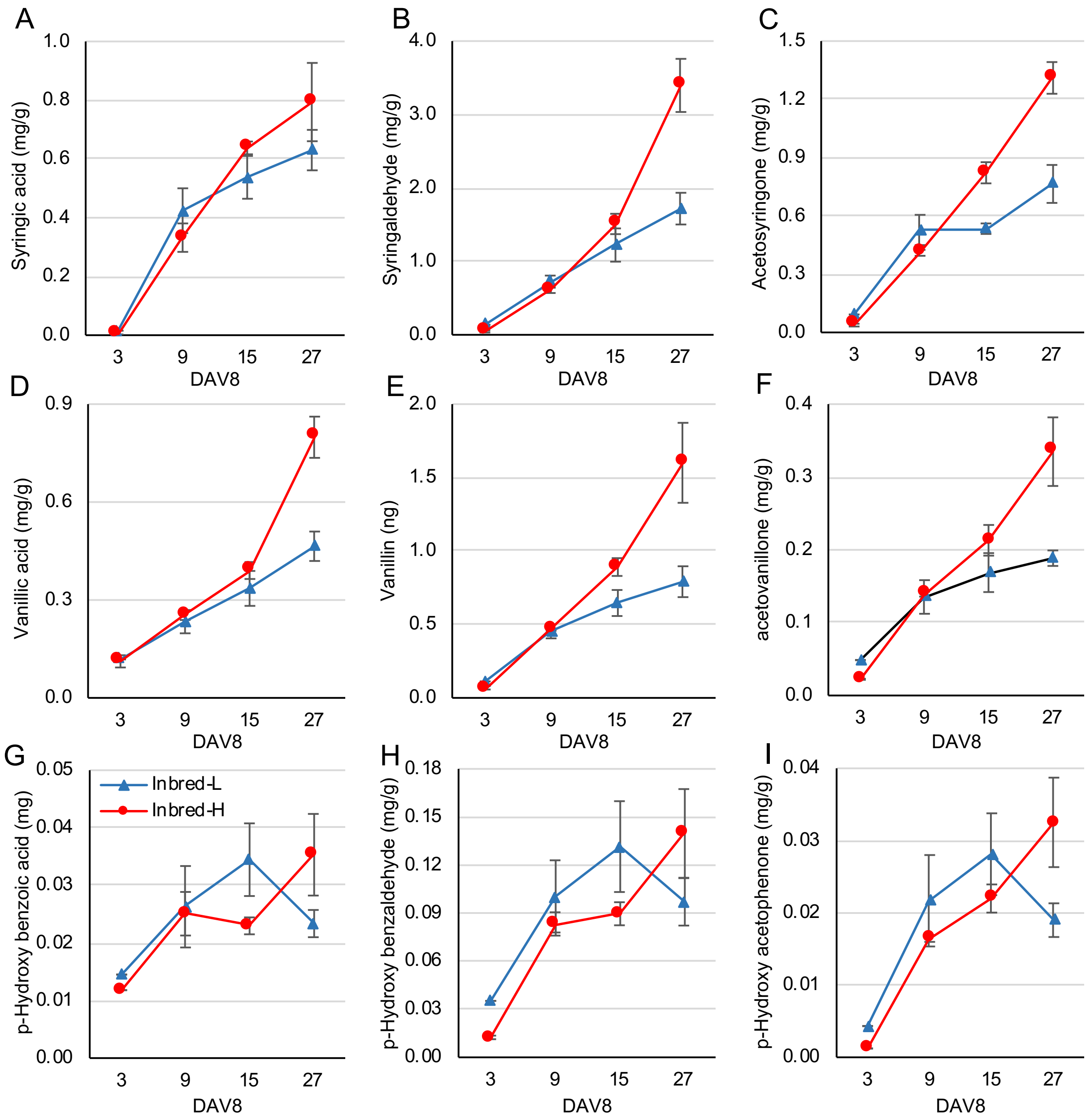


**Figure S1**. Individual monolignols constituents in internodes of Lrpr1 and Hrpr1 inbreds. Shown here are syringic acid, syringaldehyde and acetosyringone of S lignin (A-C); and vanillic acid, vanillin and acetovanillone of V lignin (D-F); p-hydroxy benzoic acid, p-hydroxy benzaldehyde, and p-hydroxy acetophenone of H lignin (G-I).
