## Supplementary material for "Genetic Architecture of Maize Rind Strength Revealed by the Analysis of Divergently Selected Populations": Figure S2

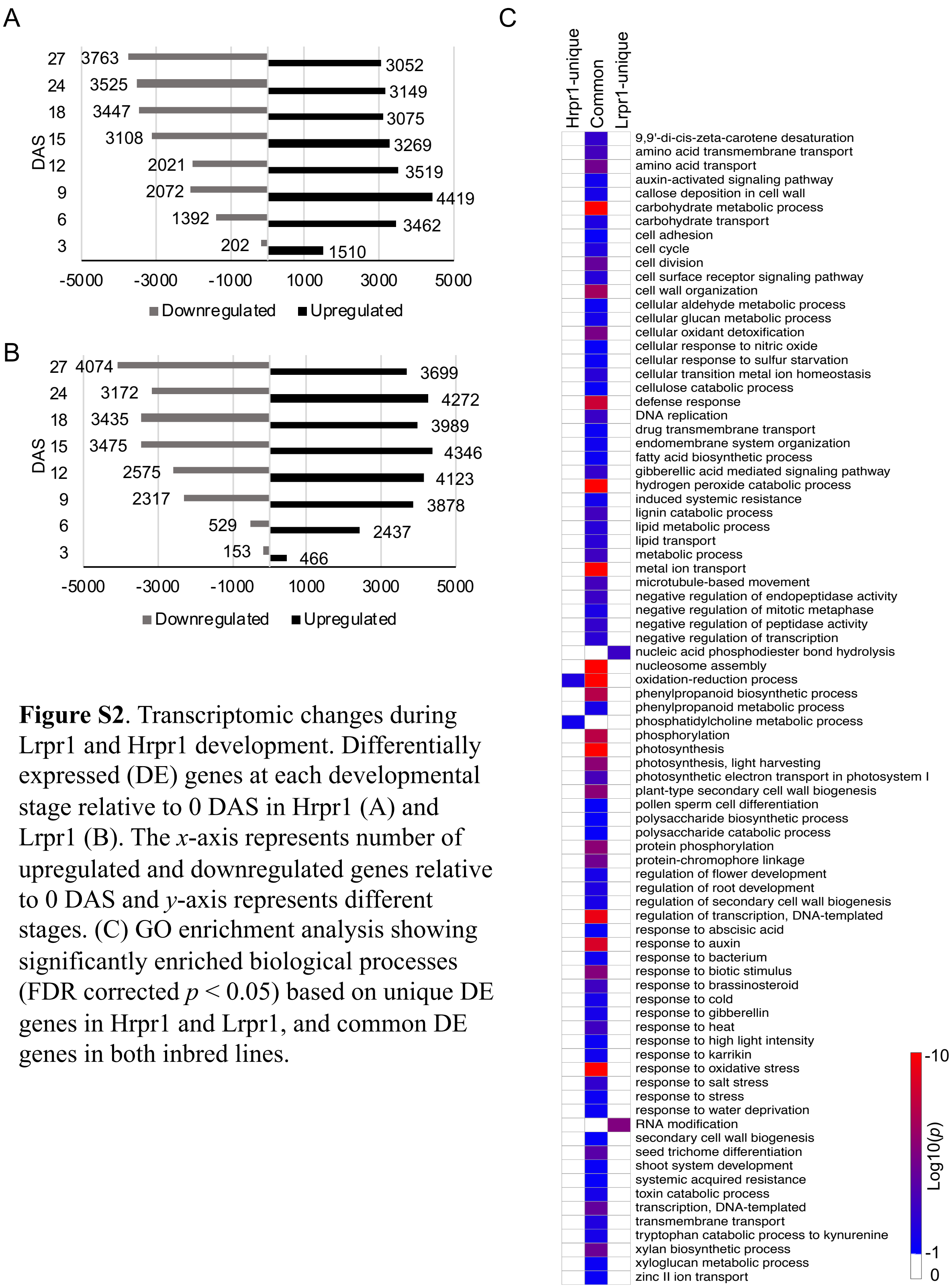


**Figure S2**. Transcriptomic changes during Lrpr1 and Hrpr1 development. Differentially expressed (DE) genes at each developmental stage relative to 0 DAS in Hrpr1 (A) and Lrpr1 (B). The *x*-axis represents number of upregulated and downregulated genes relative to 0 DAS and *y*-axis represents different stages. (C) GO enrichment analysis showing significantly enriched biological processes (FDR corrected $p < 0.05$) based on unique DE genes in Hrpr1 and Lrpr1, and common DE genes in both inbred lines.
